# TGIF2 is a major regulator of neural stem cell fate and neurogenic priming

**DOI:** 10.1101/2025.02.13.635953

**Authors:** Yiling Li, Anthi C. Krontira, Franziska Vierl, Maria L. Richter, Weixu Wang, Juliane Merl-Pham, Fabian J. Theis, Stefanie M. Hauck, Magdalena Götz

**Affiliations:** Institute for Stem Cell Research, Helmholtz Center Munich, Biomedical Center, Planegg-Martinsried, Germany; Biomedical Center Munich, Physiological Genomics, Biomedical Center, LMU Munich, Planegg-Martinsried, Germany; Graduate School of Systemic Neurosciences, LMU Munich, Planegg-Martinsried, Germany; Institute of Computational Biology, Computational Health Center, Helmholtz Munich, Munich, Germany; TUM School of Life Sciences Weihenstephan, Technical University of Munich, Germany; Research Unit Protein Science and Metabolomics and Proteomics Core, Helmholtz Center Munich, Neuherberg, Germany; School of Computation, Information and Technology, Technical University of Munich, Munich, Germany; Excellence Cluster of Systems Neurology (SyNergy), Biomedical Center, LMU Munich, Planegg-Martinsried, Germany

## Abstract

During brain development, neural stem cells (NSCs) must balance self-renewal with differentiation and ensure lineage progression. To identify novel regulators of NSCs during neurogenesis, we isolated NSCs by FACS from the mouse cerebral cortex and ganglionic eminence at mid-neurogenesis, and at birth, when gliogenesis starts in both, but neurogenesis only continues in the latter region. RNA-seq and ATAC-seq revealed major transcriptional and chromatin changes between these stages and identified TGFB-Induced Homeobox Factor 2 (TGIF2) as a key candidate factor in neurogenic NSCs. *In vitro* and *in vivo* experiments demonstrated a potent role of TGIF2 controlling NSC fate maintenance mediated by its interaction with the SIN3A/HDAC repressor complex suppressing neuronal differentiation genes. Multiomic comparison of NSC and neuron gene expression allowed the comprehensive analysis of neurogenic priming in cortical NSCs, identifying TGIF2 as its major regulator by restraining neuronal differentiation gene activation in NSCs.

## Introduction

Stem cells need to balance self-renewal versus generation of differentiated progeny during organogenesis. In the context of brain development, this balance is crucial for regulating brain size and ensuring proper neural function. During neurogenesis, neural stem cells (NSCs) are endowed with the capacity to generate neurons, while they lose this property and disappear in most brain regions, when gliogenesis starts^1^. However, in certain regions such as the murine lateral ganglionic eminence (LGE), adult neural stem cells emerge that continue to generate a subset of neurons—the olfactory bulb interneurons in mice—throughout life. This prompts two main questions: first, which factors regulate the neurogenic fate of NSCs, and second, how do NSCs generate neurons, while remaining undifferentiated themselves?

Lineage priming has been proposed in several stem cell types as a mechanism to ensure generation of the right type of progeny. In hematopoietic stem cells (HSCs) for example, the opening of regulatory elements for lymphoid genes, while keeping their expression levels low, biases HSCs toward generating the lymphoid lineage rather than other types of progeny^2^. Similarly, NSCs are primed for the generation of specific neuronal subtypes^3^, but the mechanisms retaining expression of these neuronal subtype genes at basal levels remain poorly understood^4^. However, balancing NSC fate with differentiation is essential for the timing of neurogenesis and the brain size. Thus, while significant progress has been made in understanding the transcriptional regulators of neurogenesis and gliogenesis^1,5,6^, our knowledge remains limited regarding the key factors governing neurogenic priming and equipping NSCs with neurogenic potential, while preventing their premature differentiation. Moreover, our knowledge about pan-neurogenic regulators is still rather limited^7^. Most known major potent regulators of neurogenesis, such as the proneural factors NEUROG1/2 and ASCL1, or PAX6, DLXs, and ISLET^8–11^, are expressed and function in a region-specific manner, contributing to the generation of different neuronal subtypes. However, we still know very little about non-patterned pan-neurogenic regulators.

To identify such factors, we choose to isolate NSCs labelled for CD133/Prominin1, and neurons labelled for PSA-NCAM using fluorescence-activated cell sorting (FACS) as described before^12,13^ from the cerebral cortex and the LGE. NSCs and neurons were collected at the peak of neurogenesis at embryonic day 14 (E14), and the transition to gliogenesis at E18. In the cerebral cortex, neurogenesis largely ends at E18, whereas it continues in the LGE for at least some neuronal subtypes, such as olfactory bulb interneurons, alongside the initiation of gliogenesis. This comprehensive analysis not only provides a rich resource for the first time directly comparing the two regions across developmental stages, but also led to the discovery of TGIF2 as a pan-regional key regulator of NSC fate and neurogenic priming, preventing premature neuronal differentiation and thereby maintaining the pool of NSCs during neurogenesis.

## Results

### Transcriptional and chromatin regulators of neurogenesis across forebrain regions and time

To uncover critical regulators of NSC fate and neurogenesis, we chose to perform bulk RNA-seq and ATAC-seq for deeper sequencing and more sensitive analysis of the NSCs populations isolated by FACS from E14 and E18 murine cerebral cortex and LGE (Figure 1A). Region of origin and developmental stage explained more than 90% of the variance, as seen with principal component analysis (PCA) for both the RNA-seq (Figure S1A) and the ATAC-seq data (Figure S2A), while technical aspects of the experimental procedures did not affect the distribution of the data (Figure S1B, S2B).

**Figure 1.**
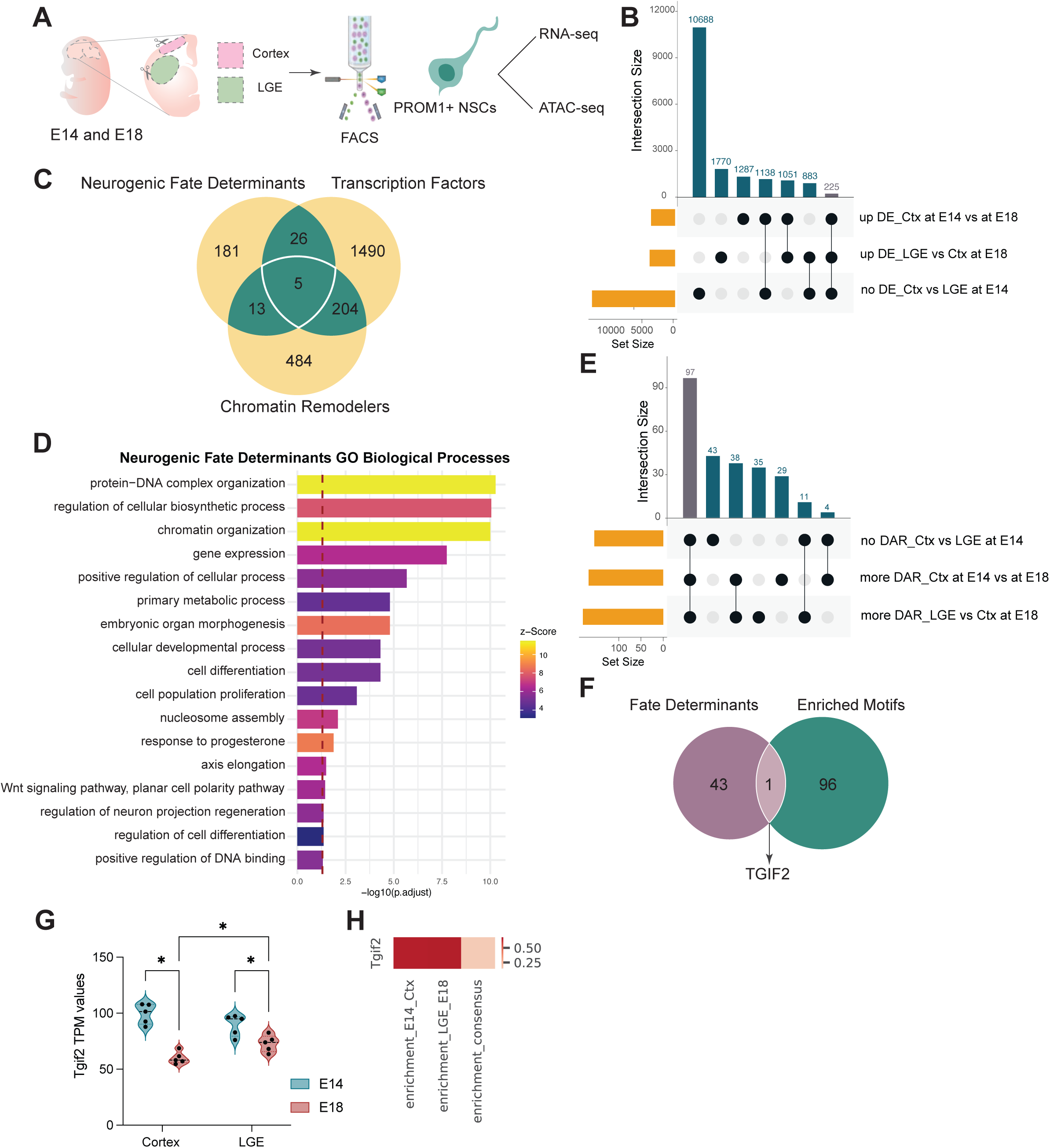
Bulk RNA- and ATAC-seq of embryonic cortex and LGE at E14 and E18. (A) Experimental scheme of RNA-seq and ATAC-seq. E: Embryonic; PROM1+: PROMININ1 (B) UpSet plot of differentially expressed genes as indicated below the plot. DE: differentially expressed; Ctx: Cortex; LGE: Lateral Ganglionic Eminence (C) Venn diagram of neurogenic fate determinants, transcription factors and chromatin remodelers. (D) GO terms associated with biological processes, showing top 2 terms each from 10 clusters of semantic space, taken from genes enriched in the neurogenic fate determinants that are transcription factors and/or chromatin remodelers. (E) UpSet plot of differentially enriched motifs at E14 versus E18 cortex, at E18 LGE versus cortex and not enriched at E14 between the regions. (F) Venn diagram of neurogenic fate determinants identified from the transcriptome analysis and the differentially enriched motifs identified in panel E. Five of the 44 neurogenic fate determinants have known binding motifs. These are *Atf3, Etv6, Mafk, Mycn* and *Tgif2*. (G) TGIF2 motif enrichment in the E14 cortex, E18 LGE and the consensus open regions shared between cortex and LGE at E14. Color represents enrichment against genomic background. (H) Tgif2 expression at E14 and E18, cortex and LGE. Significance was tested with two-way ANOVA with Benjamini, Krieger and Yekutieli correction.

Towards a comprehensive understanding of molecular regulators of neurogenesis, we first compared the transcriptome of cortical NSCs at E14 and E18 to identify transcripts with high expression at the peak of neurogenesis (E14). We found 7,455 differentially expressed (DE) genes in the cortex (at 1% false discovery rate, FDR; Table S1, Figure S1C), representing 33,55% of all detected genes (22,215 total). In the LGE, likewise 8,517 genes (38,33% of detected genes) were differentially expressed between the peak of neurogenesis and the onset of gliogenesis (1% FDR; Table S2, Figure S1D). Also the chromatin state was extensively regulated between the two developmental stages with 7,654 differentially accessible regions (DARs) in the cortex, albeit proportionally less than observed for RNA (16.2% of total) (Table S3; Figure S2C). Notably, this chromatin remodeling was less pronounced in the LGE with 4,710 accessible regions (11% of total) regulated between E14 and E18 (Table S4, Figure S2D), which may be related to neurogenesis not ending in this region. Intriguingly, a greater number of chromatin-associated factors are down-regulated at the end of neurogenesis (1% FDR: 308 for the cortex and 314 for the LGE, with 246 in common, Table S5) than up-regulated for gliogenesis (1% FDR: 52 for the cortex and 68 for the LGE, with 33 in common, Table S6). This may be consistent with larger plasticity of the neurogenic NSCs than the gliogenic NSCs. For example, we found a switch in the ATPase of the SWI/SNF complex from *Brg1* at E14 to *Brm* at E18, consistent with other tissues, where this switch occurs in more differentiated stages^14,15^.

To further explore the genes involved in the developmental stage switch we performed gene ontology (GO) term analysis and found genes with higher expression at E14 in both cortex and LGE enriched in biological pathways associated with stem cell population maintenance and differentiation, cell cycle, cell division and DNA replication (Figure S1E and S1F; Tables S7 and S8). Likewise, DARs at E14 in both cortex and LGE, compared to those at E18, were enriched with GO terms associated with nervous system development, cell differentiation and neurogenesis (Figure S2, Tables S9, and S10), further highlighting the higher degree of neurogenesis and proliferation in both regions at E14. In addition, the DE genes that were higher at E14 were enriched for nuclear localization and molecular functions such as nucleic acid and histone binding (Figure S1E and S1F; Table S3 and S4), pointing to an active role of nuclear proteins and chromatin regulation in peak neurogenesis. Thus, we specifically scrutinized transcription factors (TFs) and chromatin regulators (ChRs), as they are pivotal in regulating developmental decisions at the molecular and cellular level^12,16,17^.

We reasoned that the factors which define long-term maintenance of neurogenic stemness, thus representing essential regulators of neurogenesis, would exhibit higher expression in the E14 neurogenic cortex compared to the E18 gliogenic cortex, but would also be differentially upregulated in the E18 LGE that continues neurogenesis at larger scale compared to the cortex at the same stage. However, LGE and cerebral cortex also differ profoundly in their regional specification, as they express different patterning TFs. As we wanted to search for pan-neurogenic NSC factors, we excluded the factors that are already differentially expressed between the regions at E14 to avoid known region-specific regulators of neurogenesis. Following this rationale, we found 225 transcripts that we consider neurogenic fate regulators (Figure 1B, Table S11), 44 of which are TFs and/or ChRs (Figure 1C, Table S11). These were enriched for terms associated with regulation of developmental processes, cell differentiation and cell population proliferation (Figure 1D, Table S12), supporting our approach.

To further identify the most relevant of these TFs regulating neurogenesis, we explored which of them would have significantly more open target sites in neurogenic NSCs at E14. Following the same reasoning as for the transcriptome, we compared the differentially enriched motifs of the E14 versus E18 cortex, the E18 LGE versus E18 cortex, and focused on the ones in the commonly accessible regions between the cortex and LGE at E14. These comparisons resulted in 98 differentially enriched motifs in the neurogenic NSCs (Figure 1E, Table S13). Overlapping the 44 neurogenic fate determinants from the transcriptome analysis with the 98 neurogenic enriched motifs identified a single common key TF, whose expression and binding motifs are significantly enriched in neurogenic NSCs, namely TGFB-induced Factor Homeobox 2 (TGIF2) (Figure 1F-H). Our interest in this candidate factor was further supported by our finding that direct neuronal reprogramming of astrocytes by Neurogenin2 increases the accessibility of TGIF1 and TGIF2 binding motifs^18^, implying these factors may have a pan-neurogenic role.

TGIF2 has so far mostly been studied in the context of cancer, where it is involved in regulating migration and epithelial to mesenchymal transition^19,20^. In development, TGIF2 is a key regulator of patterning and fate in the endoderm^21,22^, but its role in the developing nervous system and neurogenesis remains unexplored. The other family member, TGIF1, is important in very early brain development where TGF-β^23^ and Sonic Hedgehog signaling regulate gastrulation and formation of the telencephalic hemispheres^24^, respectively. When TGIF1 is mutated or deleted, it can cause holoprosencephaly^24^, a phenotype where the telencephalic hemispheres are fused. Only in the adult brain, TGIF2 has been implicated in regulating behavioral aspects of autism spectrum disorder (ASD) in neurons^25^, which were improved upon TGIF2 overexpression. Thus, nothing is known about the role of TGIF2 in neurogenesis, which we decided to focus on.

### TGIF2 promotes NSC and later NPC fate in a cell-autonomous manner in vitro

TGIF2 is highly enriched in the ventricular zone (VZ), where neural stem and progenitor cells reside (Figure S3A). It has two protein-coding isoforms in rodents^26^, with the longer isoform (TGIF2IR), which contains a retained intron, being the canonical and more highly expressed isoform^26^(Figure 2A). To explore first the function of endogenous TGIF2, we started by performing knockdown experiments (TGIF2 KD) using an siRNA pool targeting all TGIF2 isoforms (Figure S3B and S3C).

**Figure 2.**
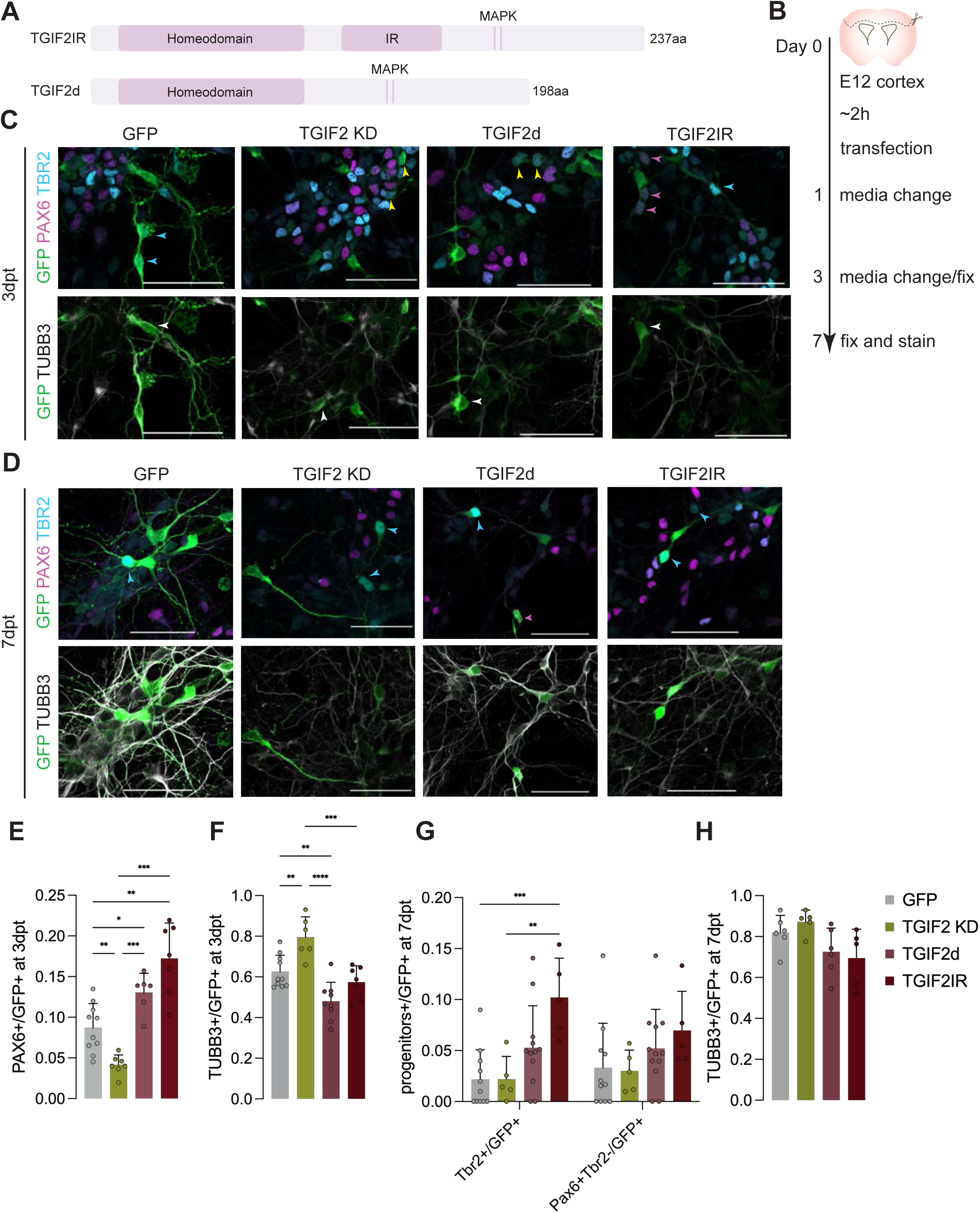
TGIF2 overexpression promotes NSC state while TGIF2 knockdown promotes differentiation. (A) Schematic drawing of TGIF2 isoforms. IR: intron retention; d: deleted; SID: SIN3a-interacting domain; MAPK: mitogen-activated protein kinase sites. (B) Schematic drawing showing the procedure of E12 cortex cells transfection assay. (C-D) Representative images showing transfected E12 cortex cell culture at 3 days or 7 days post transfection (dpt), respectively. Magenta arrowheads for PAX6+TBR2-/GFP+ cells, blue arrowheads for TBR2+PAX6-/GFP+ cells, yellow arrowheads for PAX6+TBR2+/GFP+ double positive cells. Scale bar: 50 μm. (E-F) Quantifications of PAX6+/GFP+, TUBB3+/GFP+ at 3dpt, mean±SD. N = 6-10 pools of embryos. Brown-Forsythe and Welch ANOVA tests with Dunnett’s T3 multiple comparisons test in (D), ordinary one-way ANOVA with Tukey’s multiple comparisons test in (E). * p<0.05, ** p< 0.01, *** p<0.001, ****p<0.0001. (G-H) Quantifications of TBR2+/GFP+, PAX6+TBR2-/GFP+, and TUBB3+/GFP+ cells at 7dpt, mean±SD. N = 5-12 pools of embryos. Ordinary two-way ANOVA with Tukey’s multiple comparisons test in (G) and ordinary one-way ANOVA with Tukey’s multiple comparisons test in (H). ** p< 0.01, *** p<0.001.

Cells dissociated from cerebral cortices at E12 (Figure 2B) were co-transfected with the siRNA pool and a GFP control plasmid to label proliferating cells and their progeny. At three days post-transfection (3dpt), the cells were fixed and stained for GFP, PAX6 for NSCs, TBR2 for neural progenitor cells (NPCs), and TUBB3 (tubulin beta 3 class III) for young neurons (Figure 2C and 2D). Interestingly, TGIF2 KD showed significantly reduced proportions of PAX6+ NSCs (Figure 2E) and increased proportions of TUBB3+ neurons at 3dpt (Figure 2F), suggesting a role of TGIF2 in inhibiting neuronal differentiation and favoring NSC fate.

To explore these findings further and to understand the role of the different TGIF2 isoforms, we cloned each isoform into a bicistronic expression vector driven by the CAG promoter. The vector also included GFP connected by an internal ribosome entry site (IRES) to ensure the co-expression of TGIF2 and GFP within the same cells. A monocistronic vector expressing only GFP served as the control. The constructs were transfected into dissociated cells and analyzed at 3dpt as described above (Figure 2C and 2D). Notably, we found the opposite phenotype as in the KD conditions, namely a significant increase of PAX6+ NSCs at 3dpt of both TGIF2 isoforms, with TGIF2IR showing a stronger effect (Figure 2E). Correspondingly, TUBB3+ neurons significantly decreased with the TGIF2d isoform (Figure 2F). At a later stage (7dpt), PAX6+ NSC numbers were no longer increased; however, TBR2+ NPCs significantly increased upon overexpression of TGIF2IR, with a similar but less pronounced trend for TGIF2d (Figure 2G). However, no significant difference was observed anymore for TUBB3+ neurons at 7dpt (Figure 2H), suggesting that TGIF2 overexpression promotes NSCs and delays neuronal differentiation, but does not block it. This is also consistent with TGIF2 promoting NSC maintenance initially (3dpt), followed by an enhancement of NPC fate at 7dpt.

### TGIF2 overexpression in vivo increased neural stem and progenitor cells

To probe the function of TGIF2 and their different isoforms *in vivo*, the same overexpression (OE) constructs were *in utero* electroporated (IUE) into the mouse cortex at E13 (Figure 3A). Three days post-IUE, we examined NSCs by immunostaining, using PAX6 for labelling NSCs and TBR2 for labelling NPCs (Figure 3B-D). Importantly, both TGIF2 constructs resulted in a significant enrichment of PAX6+ NSCs, with no differences observed for TBR2+ cells (Figure 3E), mirroring the effect observed at 3dpt *in vitro* (Figure 2E). As in the control condition, most PAX6+ cells after TGIF2 OE were located in the ventricular zone, corresponding to bin1 when the cortical column is divided into 5 bins and no ectopic PAX6+ or TBR2+ cells were detected in the OE conditions. Immunostaining for the mitotic protein phospho-histone 3 (pH3) revealed a more than two-fold increase in pH3+/GFP+ cells under TGIF2 OE compared to the control, with the TGIF2d isoform showing the stronger, significant effect (Figure S3D, E). These data suggest that both TGIF2 isoforms promote and prolong the NSC state.

**Figure 3.**
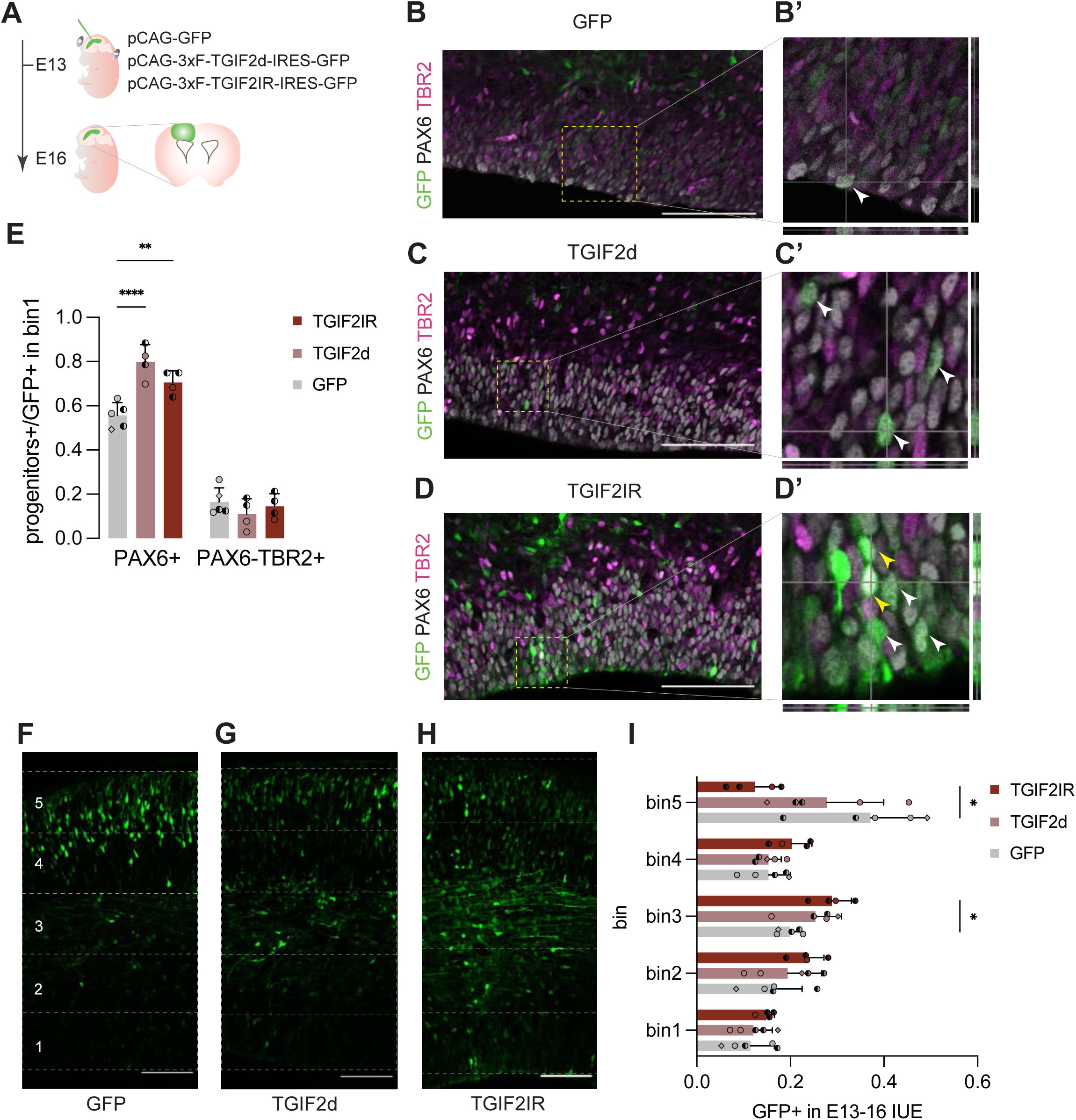
TGIF2 overexpression *in vivo* retains less differentiated cells. (A) Experimental scheme of IUE, including used plasmids. F: flag (B-D) Sections of electroporated cortices stained with PAX6 and TBR2, of which insets are to show large magnifications with orthogonal views in (B’-D’). Scale bar: 100μm. (E) Quantification of PAX6+/GFP+ and PAX6-TBR2+/GFP+ cells in bin1, mean±SD. N = 4-5 embryos from at least two different mothers. Different symbols indicate different mothers. Ordinary two-way ANOVA with Tukey’s multiple comparisons test. ** p<0.01, **** p<0.001. (F-H) Representative images of cortex 3 days after electroporation with each condition in GFP. Scale bar: 100μm. Dashed lines indicate the 5 equal bins. (I) Quantification of GFP+ cell distribution at 3 days post electroporation, mean±SD. N = 4-5 embryos from at least two different mothers. Different symbols indicate different mothers. Multiple unpaired t-tests with 5% FDR. *q<0.05

To check if the electroporated cells are stuck as NSCs or can still differentiate and migrate, the percent of GFP+ cells in each bin was determined (Figure 3F-I). We noted a trend of increased cell proportions in bins 1 and 2 upon TGIF2IR OE (Figure 3I), consistent with the significant increase in NSCs described above. The shorter isoform, TGIF2d, exhibited similar but generally milder phenotype (Figure 3I). Furthermore, overexpression of TGIF2IR resulted in a significant increase of GFP+ cells in bin 3, which contained mostly NEUROD2+ young neurons (Figure S3F-I). We also observed a concomitant reduction of GFP+ cells (by 24.7%) in the outer most bin 5, where differentiated neurons form the cortical plate (Figure 3H, I). Interestingly, the 2 isoforms differed in the size of the effect, with TGIF2d affecting proliferation stronger and TGIF2IR affecting neuronal differentiation and positioning more. However, both isoforms prolonged the NSC state leading to an increase of immature neurons in bin3 and a decrease of mature neurons in the cortical plate (bin5), highly reminiscent of the *in vitro* phenotype with reduced neuronal numbers.

### TGIF2 overexpression reduces cells expressing more mature neuronal differentiation genes shown by scRNA-seq

To investigate transcriptomic changes underlying TGIF2’s effect in retaining NSC state, we performed scRNA-seq on GFP+ cells isolated 36 hours post-IUE using FACS (Figure 4A). A total of 51,392 cells were obtained after quality control filtering. Dimensionality reduction via UMAP showed consistent overlap among conditions and replicates (Figure S4A). Cell clusters were identified via the Leiden algorithm (Figure S4B) and annotated based on marker gene expression (Figure 4B, Figure S4D, E). For instance, we identified NSCs, marked by *Pax6*, *Sox2*, and the radial glia marker *Fabp7* (Fatty acid binding protein 7), and NPCs, marked by expression of *Eomes* (also known as TBR2), *Neurog2*, and *Elavl2* (Figure S5E). Cell cycle phases were inferred through cell cycle marker genes to identify cycling cell populations (Figure S4C).

**Figure 4.**
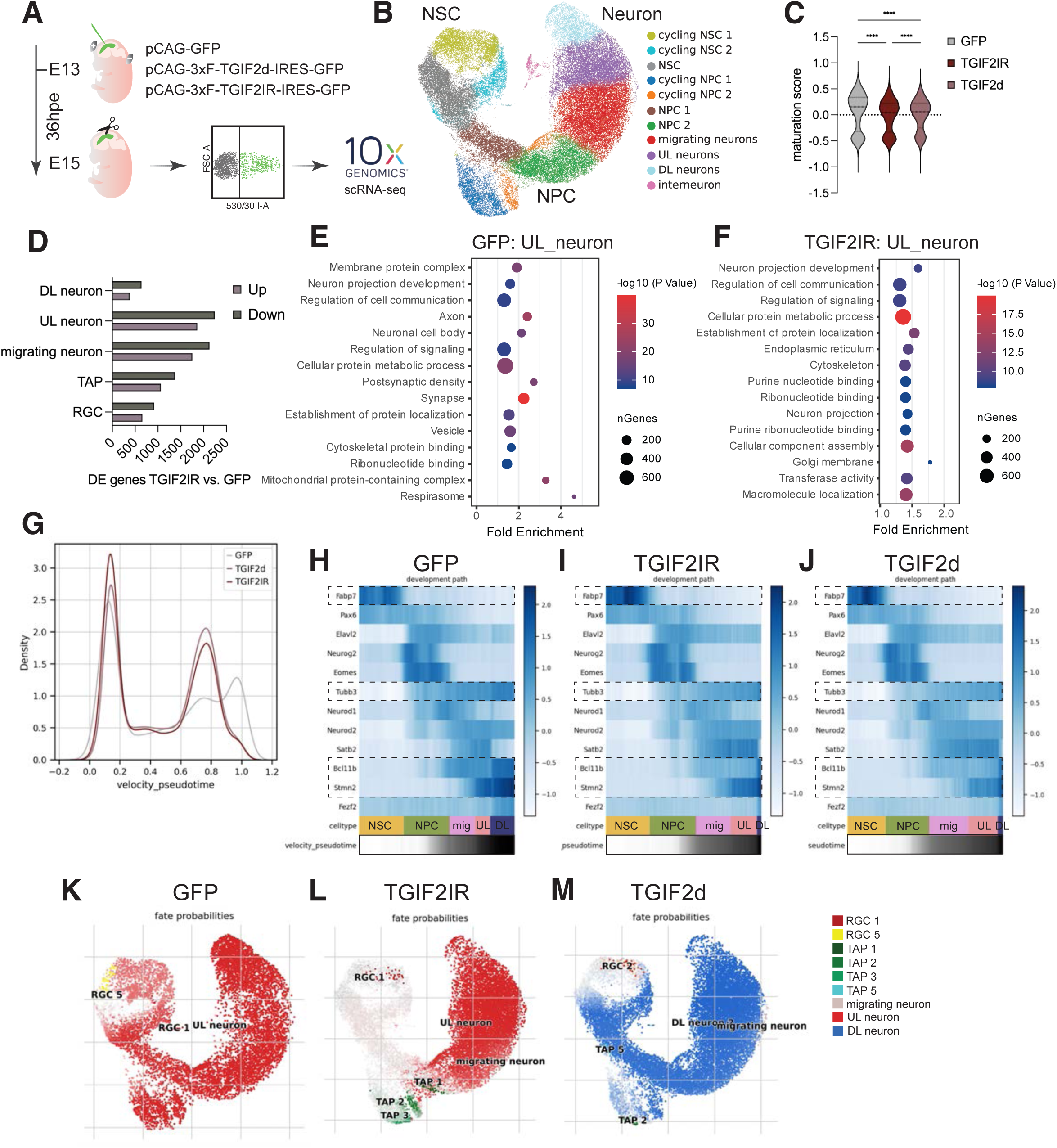
TGIF2 overexpression slows differentiation shown by scRNA-seq. (A) Schematic drawing of experimental procedures. (B) UMAP projection with each cluster annotated with corresponding cell type. (C) Violin plot of maturation score per condition. Kruskal-Wallis test with Dunn’s multiple comparisons test. (D) Barplot of DE genes between TGIF2IR and GFP identified in each cell cluster. (E-F) Top 15 terms of GO term enrichment analysis of DE genes in upper layer neurons between GFP and TGIF2IR. (G) Cell density plot along velocity pseudotime. (H-J) Gene expression of selected markers by velocity pseudotime and differentiation lineage. (K-M) Fate probability maps from CellRank^28,29^ analysis. RGC: radial glial cells, TAP: transit-amplifying progenitors, DL neurons: deep layer neurons, UL neurons: upper layer neurons.

Comparing TGIF2 expression levels between GFP control (representing endogenous TGIF2 levels) and TGIF2IR conditions revealed that TGIF2IR overexpression was prominent in all stem and progenitor clusters, migrating neurons and in upper layer neurons (UL neurons) (Figure S4F). Using a maturation score (Figure 4C), calculated by average expression of genes related to neuronal maturation (see Methods), we could observe that cells were in a less mature state in TGIF2 conditions, with TGIF2IR being even less mature than TGIF2d. We then conducted DE analysis across cell types, which revealed that more genes were downregulated in TGIF2IR over-expression condition compared to the control (Figure 4D), highlighting a potential repressive role of TGIF2. GO analysis on DE genes within UL_neuron (upper layer neuron) cluster showed general terms for neurogenesis, such as “neuron projection development” and “regulation of cell communication” for both GFP control and TGIF2IR conditions (Figure 4E, F), but TGIF2IR did not acquire terms for a more mature state, such as “axon” and “postsynaptic density” (Figure 4E). Altogether, these data suggest that TGIF2-overexpressing conditions result in transcriptomic downregulation across all cell types, which are in a less mature state of differentiation. This maturation difference prompted us to conduct pseudotime trajectory analysis.

RNA velocity pseudotime analysis, based on spliced and unspliced RNA ratio^27^, uncovered that TGIF2IR-expressing cells remained predominantly in early differentiation stages, while GFP control cells having progressed to later stages of differentiation (Figure 4G, Figures S5A-C). This delayed differentiation across pseudotime is particularly evident in NPC 2 and post-mitotic neurons (Figure S5D-F). This is concomitant with higher expression of *Fabp7* in NSCs and lower expression of neuronal genes (*Tubb3, Bcl11b, Stmn2*) in neuronal clusters in the TGIF2IR condition (Figure 4H-J). Additionally, CellRank analysis^28,29^ based on RNA velocity assigned 13 macrostates for fate prediction (Figures S5G-R). Interestingly, TGIF2IR overexpression delayed the assignment of upper layer neuron fates and maintained cells more in NPC states (Figures 4K and 4L). The overexpression of the shorter isoform TGIF2d also delayed differentiation, maintaining some cells in NPC states, although the effect was slightly less pronounced than with TGIF2IR (Figure 4M). Surprisingly, there was no upper layer neuronal fate being predicted in TGIF2d condition, but only deep layer neuronal (DL neuron) fate (Figure 4M, Figure S5O-R). Collectively, these analyses unbiasedly confirmed that TGIF2 overexpression downregulates expression of neuronal differentiation genes, and upregulates genes in NSCs, thereby maintaining cells in progenitor states, similar to our findings based on immunostainings (Figures 2J, 3D-F).

### TGIF2 binds and negatively regulates neuronal differentiation genes

To better understand the molecular mechanisms by which TGIF2 TFs promote NSC fate and limit neuronal differentiation, we first aimed to identify direct binding targets, focusing on TGIF2IR, as it is the major isoform expressed and generally had stronger effects *in vivo*. Cut&Run analysis was performed after dissecting IUE regions based on GFP at 36 hours post-electroporation, the same timepoint as the scRNA-seq (Figure 5A). This analysis uncovered 10,688 peaks (Figure 5B), with TGIF2IR predominantly binding to intronic (44.57%) and intergenic regions (33.5%), indicating a preference for proximal regulatory elements over promoters (10.23%) (Figure 5C). Motif enrichment analysis identified many important TFs for neurogenesis and neuronal differentiation, such as ASCL1, NEUROD2, NEUROG2, MEIS1,2, and MYC (Figure 5D). Although TGIF2 itself was not among the top enriched motifs, motif scanning analysis using the known TGIF2 motif identified 3,176 occurrences (p-value <0.001) among the peaks (Table S14). It is worth noting that the known TGIF2 motif was derived from ChIP-seq data in mouse embryonic stem cells, which may differ from the motif in neural stem and progenitor cells.

**Figure 5.**
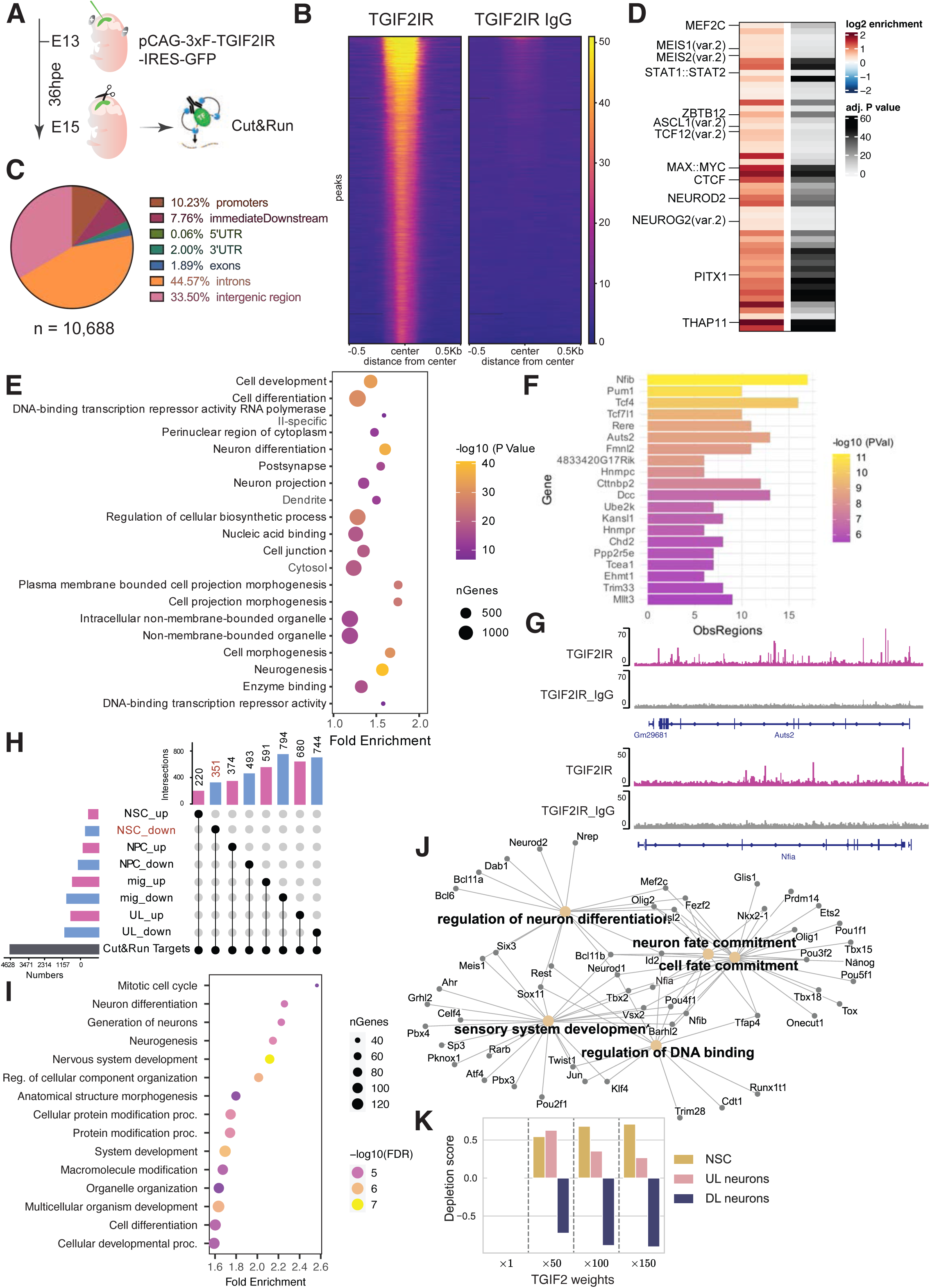
TGIF2 binds at neural differentiation genes and regulates chromatin factors. (A) Schematic drawing of experimental procedures of Cut&Run. (B) Enrichment heatmap of TGIF2IR peaks and its corresponding IgG control, centered at the middle of the peaks. (C) Pie chart of genomic distribution of TGIF2IR peaks. (D) MonaLisa motif enrichment analysis of TGIF2IR peaks. (E) Top 20 terms from GO term enrichment analysis of annotated genes. (F) Top 20 genes with extensive regulation by GREAT. (G) Peak examples with bigwig profiles exported from IGV^86^. (H) UpSet plot overlapping Cut&Run targets and DEGs in scRNA-seq between TGIF2IR and GFP per cell type. (I) Top 15 enriched GO terms in biological processes of overlaps (351 genes) between TGIF2IR Cut&Run targets and downregulated genes in NSCs of TGIF2IR compared to GFP control from scRNA-seq. (J) GRN built by CellOracle^79^, representing negatively regulated TFs by TGIF2 and associated GO terms. (K) Weighted simulations by RegVelo^31^ for TGIF2 overexpression effect on cell fate bias.

Annotation of the nearest genes to the identified peaks revealed 5,783 target genes (Table S15). GO analysis of these targets showed significant enrichment in terms such as “neurogenesis,” "postsynapse,” “dendrite,” and "cell projection morphogenesis," all of which are crucial processes in neurogenesis, supporting cell migration and synaptic maturation (Figure 5E). In addition, we also found TGIF2 targeting many RNA-binding and splicing factors (e.g. *Stau1/2*, *Pum1/2*, *Ptbp2*, *Snrnps*) and signaling mediators, such as *Tle4*, *Tcf7l1* and *Smad4*. Further examination of peak distribution using GREAT^30^ identified genes highly regulated by TGIF2IR (Figure 5F, Table S16). For instance, *Auts2* and *Nfia* were associated with around 20 peaks across their gene bodies, indicating extensive regulation by TGIF2IR (Figure 5G). These highly regulated genes are associated with “H4 histone acetyltransferase complex” (*Kansl1*, *Epc1*, *Mllt3*), “growth cone” (*Dcc*, *Auts2*, *Myh10*), and “chromatin” (*Brd4*, *Smarcc1*, *Arid1b*) (Table S17), highlighting TGIF2 as an upstream regulator of chromatin factors and neuronal differentiation genes.

In order to determine the importance of these direct targets, we performed two analyses: (1) overlaying them with genes regulated by TGIF2 in scRNA-seq and (2) using RegVelo^31^ to explore the transcriptional networks influenced by TGIF2. For the first analysis, we overlapped annotated genes from TGIF2 Cut&Run peaks with the DEGs between TGIF2 and GFP conditions from scRNA-seq across all cell types (Figure 5H). In general, there were more overlaps in the downregulated genes by TGIF2, constituting more than half of the DEGs in most cell types, suggesting TGIF2’s function as a transcriptional repressor. Focusing on the NSCs, the downregulated genes were enriched in GO terms such as “neuron differentiation” and “neurogenesis” (Figure 5I), the central regulated terms of TGIF2 (Figure 5E).

To understand this regulation by TGIF2 further, we used RegVelo, which relied our bulk ATAC-seq data from E14 cortical NSCs and TGIF2IR Cut&Run data for building a *priori* Gene Regulatory Network (GRN) to perform dynamic inference on scRNA-seq GFP control dataset (Figure 5J). This GRN revealed a network of targets negatively regulated by TGIF2 and highlighted “neuron fate commitment” and “neuron differentiation” as key regulated terms. Among these RegVelo-refined and negatively regulated targets, Fezf2 and Bcl11b are two critical TFs for DL neuron fate, suggesting TGIF2 may have a repressive role on DL neurons production. Indeed, when we applied weighted simulations in RegVelo to mimic TGIF2 OE, TGIF2 weights promoted NSC and UL neuron fates, simultaneously depleting DL neuron progeny (Figure 5K). Also, the more added weights we simulated, the bigger the enrichment in NSC fate, resonating with the phenotype *in vitro* and *in vivo* (Figure 2 and Figure 3).

### TGIF2 interacts with HDAC1/2 and SIN3 co-repressor complex

Seeing that TGIF2 downregulates neuronal differentiation genes and directly binds to neurogenesis associated genes, we examined if it acts as a transcriptional repressor during neurogenesis. To determine its interaction with possible repressors, we performed mass spectrometry after co-immunoprecipitation (co-IP-MS) of FLAG-tagged TGIF2IR transfected in P19 cells in two independent replicates (Figure 6A). The results (LFQ intensity ratio more than 3-fold in TGIF2IR compared to GFP; Table S18) revealed that TGIF2IR robustly interacts with components of the SIN3A co-repressor complex, including HDAC1/2^32^ and RBBP4/7, as well as lamina-associated proteins such as BANF1 and TMPO (as known as LAP2), which are known to mediate gene repression through chromatin localization^33^. Additionally, we identified interactors involved in cell cycle regulation (RPA1/2/3) and metabolism (PARP1, SSBP1) (Figure 6B). These findings confirm that TGIF2 associates with repressor proteins, specifically within the SIN3A co-repressor complex, consistent with previous data^32^ thereby further supporting its role as a transcriptional repressor.

**Figure 6.**
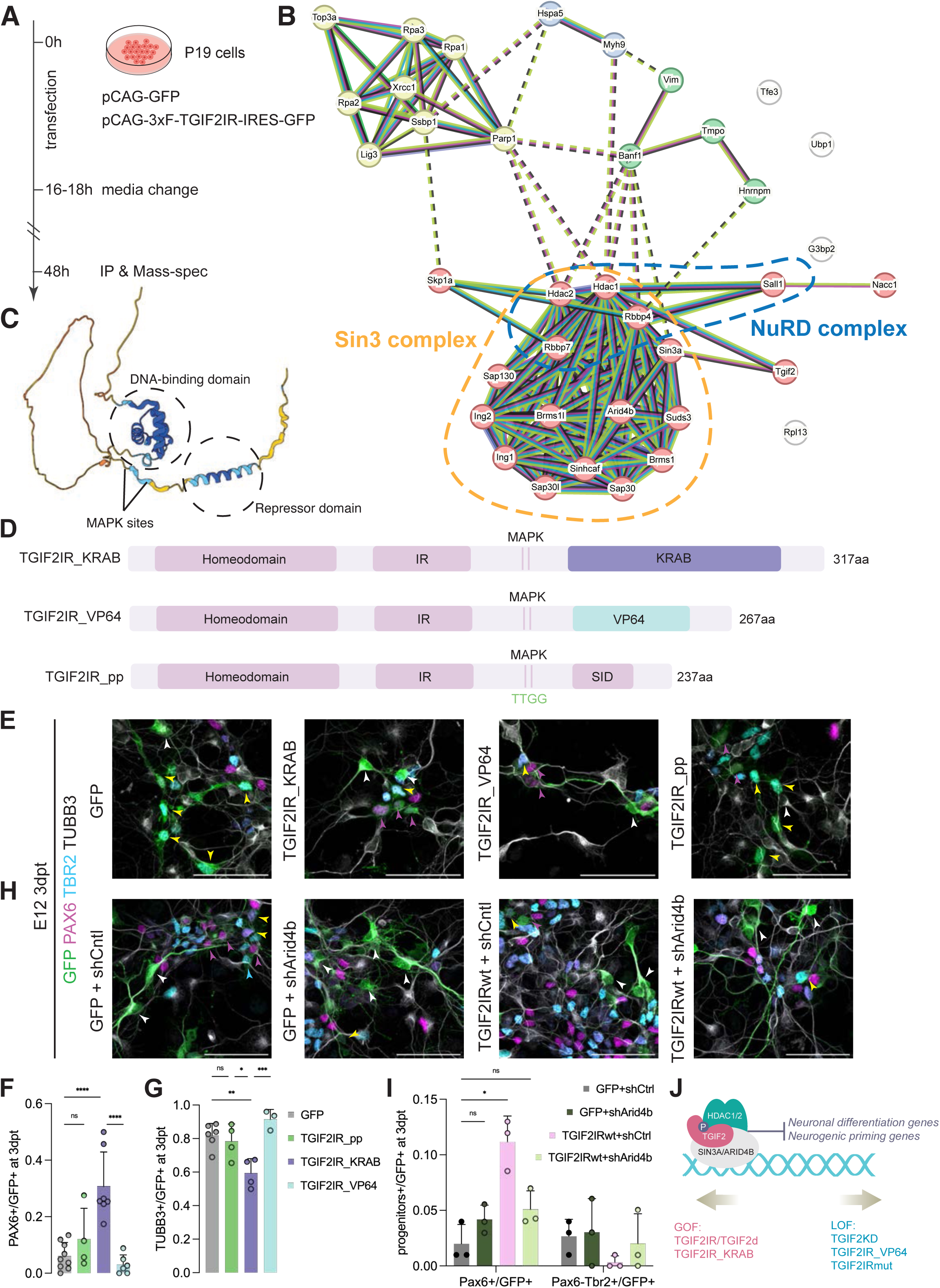
TGIF2 interacts with the SIN3a complex and acts as a repressor. (A) Schematic drawing of IP-MS experiment in P19 cells. (B) STRING analysis of interactors of TGIF2IR with LFQ intensity more than 3-fold compared to GFP control. (C) AlphaFold prediction of TGIF2 structure, with DNA-binding domain and repressor domain circled, and 2 MAPK sites indicated. (D) Schematic structures of different TGIF2 constructs. pp: phospho-mutant, TGIF2IRmut: TGIF2IR mutant form. (E, H) Representative pictures of E12 primary cortex cells cultures transfected with different conditions at 3dpt, co-stained with PAX6, TBR2, and TUBB3. Magenta arrowheads for PAX6+TBR2-/GFP+ cells, yellow arrowheads for TBR2+PAX6-/GFP+ cells, white arrowheads for TUBB3+/GFP+ cells. Scale bar: 50 μm. (F-G) Quantification of PAX6+/GFP+ and TUBB3+/GFP+ in transfected E12 culture at 3dpt with TGIF2IR_pp, TGIF2IR_KRAB and TGIF2IR_VP64 constructs, mean+SD. N = 3-9 pools of embryos. Ordinary one-way ANOVA with Tukey’s multiple comparisons test. ns: not significant. (I) Quantification of PAX6+TBR2-/GFP+ and PAX6-TBR2+/GFP+ in transfected E12 culture at 3dpt with shArid4b constructs, mean+SD. N = 3 pools of embryos. *p = 0.0341. Ordinary two-way ANOVA with Šídák’s multiple comparisons test. (J) Scheme of molecular mechanisms of TGIF2: when TGIF2 is phosphorylated, it is able to interact with SIN3A complex including ARID4B and HDAC1/2, which altogether repress neurogenesis programs, including *Arid4b* itself, to maintain NSC fate. TGIF2 KD, TGIF2IR_VP64 and TGIF2IRmut act in the opposite direction from wild type TGIF2s and TGIF2IR_KRAB.

### TGIF2 function is dependent on its repressor domain and phosphorylation

TGIF2 has been reported to exhibit repressor activity in various cell types, particularly in cancer cells^20,26^, but has also been reported to act as a co-activator^34^. To functionally manipulate repressor and activator functions of TGIF2, we first aimed to identify the repressor domain within TGIF2, utilizing sequence alignment with its paralog, TGIF1, which is better characterized^35^. This alignment revealed that the SIN3A-interacting domain (SID), interacting with the SIN3A co-repressor complex in TGIF1 and suggested to maintain pluripotency^35,36^, is conserved in TGIF2, in line with our findings in co-IP-MS.

To explore the function of the SID, we replaced it either by a more potent repressor domain, KRAB, or by an activator domain, VP64 (Figure 6D). Overexpression of TGIF2IR-KRAB in E12 dissociated cortical cell cultures resulted in an even stronger phenotype than TGIF2IR, showing a significantly higher proportion of PAX6+ NSCs (32.8%), compared to TGIF2IR (17.2%) and control (8.7%) (Figure 6E-F). This was accompanied by a substantial reduction in the neuronal population in the TGIF2IR-KRAB condition (Figure 6G). Conversely, the overexpression of TGIF2IR-VP64 led to a drastic decrease in progenitors (Figure 6E-F), with over 90% of cells differentiating into neurons (Figure 6G), thus indicating that activating TGIF2-repressed targets strongly promotes neuronal differentiation. These data suggest that TGIF2 represses neuronal differentiation genes and thereby promotes NSC fate.

Protein structure prediction of TGIF2IR using AlphaFold^37^ suggested MAPK phosphorylation sites potentially linking the DNA-binding homeodomain to the SID repressor domain (Figure 6C). Phosphorylation has been shown to regulate TGIF2 function in other contexts, particularly cancer cells^20^. To examine the role of phosphorylation of TGIF2IR in neurogenesis, we generated a phosphorylation-deficient TGIF2IR mutant by substituting the two MAPK threonine residues with glycine (TGIF2IR_pp) (Figure 6D). Overexpression of this phospho-resistant TGIF2 in E12 cortical cultures did not affect NSC maintenance or neuronal differentiation (Figure 6E-G), suggesting that TGIF2’s function in promoting NSCs is mediated by the phosphorylated form.

Collectively, these findings demonstrate that phosphorylated TGIF2 represses neurogenesis-related genes and retains NSCs and later NPCs by interacting with SIN3A co-repressor complex (Figure 6H).

### A mutation in SID overturns the effect of TGIF2 and unravels interactors essential for TGIF2 function

Given the critical role of the SID in TGIF2 function, we introduced a point mutation within SID (A210V), referred to as TGIF2IRmut (Figure S6A). Overexpression of this TGIF2IRmut in E12 cortical cell cultures lead to an increase of TUBB3+ neurons promoting differentiation (Figure S6B-D). This suggested that the mutation in the SID domain abrogates the normal repressor function of TGIF2. Indeed, the effects obtained with the TGIF2IRmut were very similar to the TGIF2 KD (Figure 2F). To explore if this is also the case *in vivo*, we employed the same IUE paradigm as described above (Figure 3A). Overexpression of TGIF2IRmut resulted in a phenotype opposite to TGIF2IRwt (Figures S6E and S6F), as more cells were found in bin5, corresponding to the cortical plate, where most mature neurons are located (Figure S6G). Indeed, morphology and immunostaining confirmed that these are neurons, especially UL neurons (Figure S6H) supporting that TGIF2IRmut OE causes faster neuronal differentiation also *in vivo*.

To get a comprehensive idea of how gene expression is changed by the TGIF2IRmut, we performed scRNA-seq and Cut&Run experiments as described above. Using Cut&Run, we observed a surprisingly large number of targets bound by TGIF2IRwt no longer detected in TGIF2IRmut (Figures S6I-J). This included *Arid4b* (Figure S6K), a component of the SIN3A complex that interacts with the SID domain of canonical TGIF2IRwt (Figure 6B). To understand how this loss of binding affects gene regulation, we overlaid genes aberrantly upregulated in TGIF2IRmut (Table S19) with the peaks bound by TGIF2IRwt, but not the TGIF2IRmut (Figure S6L). This showed an interesting signature revealing *Gatad2* as differentially bound and regulated (Table S20). This factor is part of the NURD complex that regulates neuronal activity genes^38^. In addition, mutations of *Gatad2* cause delayed neuronal differentiation in patients, highlighting *Gatad2* as a possible key down-stream effector^39^. We further found stem cell factors, such as *Vcam1* and *Fabp7* affected in their expression (Table S20), alongside with many genes involved in translation and proliferation. Thus, lack of DNA-binding and target gene regulation leads to the loss of TGIF2 function upon the mutation in the SID domain.

Next, we aimed to explore, if also the interactome of this TGIF2IRmut would differ from the TGIF2IRwt in P19 cells (Figure S6M). Interestingly, interactome changes were less abundant than those seen in Cut&Run, revealing the loss of only 7 protein interactions for the TGIF2IRmut compared to TGIF2IRwt (Figure S6N). Amongst them we observed again ARID4B. Thus, *Arid4b* is not only a direct target of TGIF2IRwt, that is no longer bound by TGIF2IRmut, but also an interactor of TGIF2IRwt.

As both the Cut&Run and interactome pointed to a key role of ARID4B involved in TGIF2 function, we examined if ARID4B is essential for the function of TGIF2IRwt. Using the same E12 assay as described above, we transfected either an shRNA targeting the open reading frame of *Arid4b* (Figure S8O), following a GFP reporter (pCAG-GFP-shArid4b), or a non-targeting control shRNA (pCAG-GFP-shCtrl), either with the GFP control (pCAG-GFP) or with TGIF2IRwt (pCAG-TGIF2IRwt-IRES-GFP) vectors. KD of *Arid4b* abolished the effect of TGIF2IRwt overexpression in retaining PAX6+ NSCs (Figure 6I), confirming our hypothesis that ARID4B interaction is necessary for TGIF2IR’s repressor function. We have thus identified a crucial interactor and down-stream target of TGIF2 involved in its key functions in neurogenesis.

### TGIF2 as major regulator of primed neuronal lineage genes in NSCs

Given the function of TGIF2 in repressing neuronal differentiation genes in NSCs, we considered that TGIF2 could be involved in lineage priming by restraining the expression of neuronal genes that may be accessible already in NSCs. In other stem cell systems, lineage priming involves the opening of regulatory elements for progeny-specific genes, while restraining their expression levels^2^. However, the mechanisms underlying neurogenic priming in NSCs are still poorly understood^3,40^. To address this, we stained for PSA-NCAM to isolate neurons by FACS from the E14 cerebral cortex (Figure 7A) and performed RNA-seq to identify differential gene expression between neurons and NSCs. Among 5835 DEGs higher in neurons than NSCs (at FDR 1%), 4984 (85.4%) displayed open chromatin accessibility in our ATAC-seq of cortical NSCs already at E14, thus fitting the definition of priming with being accessible but lower expressed than later in the lineage. Focusing on the genes whose regulatory sites experienced significantly reduced accessibility in NSCs at the end of neurogenesis (E18), we identified 433 genes, which we named as neurogenic priming genes (Figure 7B, Table S21). Notably, 225 of these genes (51.9%) were direct targets of TGIF2, as determined by our Cut&Run data (Figure 7B, Table S22). Both the neurogenic priming genes and the TGIF2-regulated subset were enriched for GO terms such as “axonogenesis” and “neuron differentiation” (Figure 7C and 7D). Our data revealed that TGIF2 binds directly to these accessible chromatin regions of priming genes in NSCs, as exemplified in Figure 7E. To assess if the enrichment of TGIF2 targets in neurogenic priming genes is significant, we generated 100,000 permutations of equal-sized, randomly selected gene sets that are not TGIF2 targets. Remarkably, no other gene set exhibited more than 77 overlapping genes with the neurogenic priming gene set (Figure 7F), underscoring the specificity and importance of TGIF2’s regulatory role on neurogenic priming. Together, these findings identify TGIF2 as not only a novel and non-patterned regulator of neurogenesis, but also a major regulator of neurogenic priming in NSCs.

**Figure 7.**
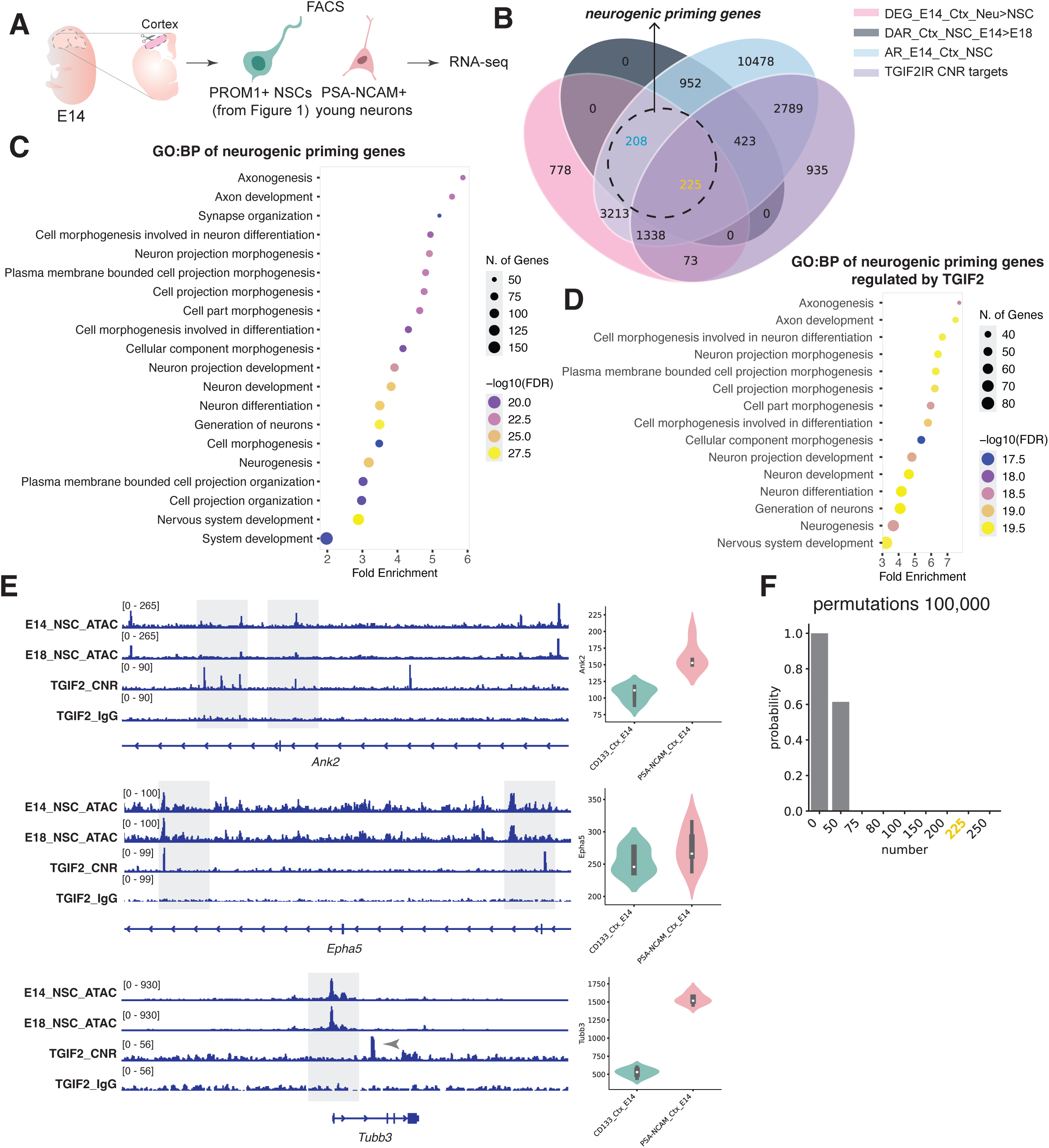
TGIF2 is a master regulator of neurogenic priming. (A) Scheme for the experimental setup: E14 cortices were dissected for FACSorting of PSA-NCAM+ young neurons and performed RNA-seq, to compare with E14 cortical NSCs RNA-seq data mentioned in Figure 1. (B) Venn diagram showing data mining logic of neurogenic priming genes. DEG: differentially expressed genes. DAR: differential accessible regions. AR: accessible regions. CNR: Cut&Run. Neu: Neurons. Ctx: Cortex. NSC: Neural Stem Cells. E: embryonic. (C) GO term analysis of neurogenic priming genes. (D) Examples of neurogenic priming genes regulated by TGIF2, showing ATAC-seq and TGIF2 Cut&Run (CNR) profiles, together with each gene’s RNA expression by violin plot in E14 cortical NSCs and neurons. (E) Permutation test with 100,000 trials to identify the possibility of various number of random gene sets to overlap with the neurogenic priming genes.

## Discussion

Here we provide a comprehensive resource profiling RNA- and ATAC-seq data in NSCs across distinct brain regions, and timepoints—at the peak of neurogenesis and onset of gliogenesis, with one region continuing neurogenesis. This dataset not only enables the identification of novel pan-neurogenic regulators, exemplified by the hundreds of candidates listed in Table S11. Among these, we focused on TGIF2, showcasing its critical role in regulating NSC fate and neurogenic priming. Beyond TGIF2, our dataset also provides insights into chromatin and epigenetic regulatory dynamics during this fundamental switch in lineage transition from neurogenesis to gliogenesis. Notably, the differential chromatin regulators between these two stages provide a valuable entry point towards a better understanding of this transition across regions. To facilitate further exploration of this dataset, a Shiny App will be made publicly available.

Our focus on TGIF2 stemmed from its expression pattern correlating with neurogenesis across regions and its enriched motif within neurogenesis-restricted open chromatin regions identified by ATAC-seq. We showed TGIF2 as a key regulator of NSC maintenance and neuronal differentiation by knock-down and overexpression experiments. TGIF2 functions as a molecular “brake” on neurogenesis programs, actively gatekeeping NSC and later NPC states, thereby interfering with premature differentiation and fine-tuning the timing of cortical development. By integrating single-cell transcriptomics, Cut&Run, proteomics, and functional assays of fusion and mutant proteins, we demonstrated that TGIF2 maintains NSC fate not through its canonical role in antagonizing TGFβ signaling^19^, but rather by repressing neuronal differentiation genes targeted by key neurogenic regulators. This function allowed us to discover that TGIF2 is a major regulator of neurogenic priming.

Lineage priming can occur via transcriptional priming, where genes defining the later lineage are already expressed in stem or progenitor cells at low mRNA levels without protein translation^3^, or via epigenetic priming, where regulatory sites of these genes are open and sometimes epigenetically “poised” or “primed” by specific marks^2^. Here we considered priming genes in NSCs as those expressed significantly higher in neurons, but already with open chromatin in NSCs at E14. Remarkably, TGIF2 bound more than half of them. As it is a pan-neurogenic factor expressed not only in LGE and cortex NSCs, but throughout CNS regions^41^, we would propose TGIF2 functions as a central regulator of neurogenic priming in a wider context. Supporting the wider relevance of our data also across species, RNA-seq data from the human cortex also revealed that TGIF2 expression steeply declines at gliogenesis stages (post-conceptual week 20)^42^. Thus, TGIF2 represents a novel regulator of neurogenesis and neurogenic priming, complementing the translational repression previously described^4^. TGIF2-mediated transcriptional repression allows primed NSCs to remain poised for differentiation cues and respond in a timely manner during the dynamic changes in neurogenesis. By maintaining basal expression levels of neuronal differentiation genes, TGIF2 ensures NSCs are primed for lineage commitment without undergoing premature differentiation.

In this regard, it is also important to mention that TGIF2 itself is regulated by signaling pathways, namely MAPK/ERK signaling induced phosphorylation, as shown before in cancer cells^20,43^. Mutating the two MAP kinase phosphorylation sites in TGIF2 completely abolished its ability to promote NSC fate. Interestingly, proteomic analysis of human iPSC-derived NSCs and neurons^44^ revealed that TGIF2 is phosphorylated only in NSCs, but not in neurons, while its total protein levels remain unchanged (data not shown). Indeed, the activation of MAPK/ERK is required for NSC proliferation, and has to decline for neuronal differentiation^45,46^. ERK activity is also suggested to be a gating mechanism for neural differentiation, as inhibition of ERK induced more accessible chromatin and precocious transcription of neural genes in spinal cord precursors^48^. These findings suggest that TGIF2’s activity is developmentally regulated by endogenous signaling pathways, such as MAPK/ERK signaling^20,43^, modulating TGIF2’s interaction with the SIN3A complex.

Notably, TGIF2 binding sites determined by Cut&Run are enriched with motifs for proneural TFs, such as ASCL1 and NEUROG2, as well as NEUROD2, MEIS1 and 2, which are known to promote neurogenesis and neuronal differentiation in both developmental as well as adult contexts^47,49–52^. This suggests that TGIF2 occupies neuronal differentiation gene loci to repress targets of and/or sterically block the access of proneural TFs, thereby inhibiting premature neural differentiation. This interplay between TGIF2 and neurogenic TFs may serve as a checkpoint to ensure the proper timing of neural differentiation during cortical development. Additionally, among the genes repressed by TGIF2, we observed significant regulation of the nuclear factor I (NFI) family of TFs, including *Nfia*, *Nfib*, and *Nfix*, which are known to function synergistically^53^. Double knockout of *Nfia* and *Nfib* has been shown to cause ventricular enlargement from progenitor proliferation and reduced neural differentiation^54^, a phenotype resembling TGIF2 overexpression–increased neural stem/progenitor cells and delayed differentiation. This finding places TGIF2 upstream of NFI family members in the regulatory hierarchy, functioning as a negative regulator of neuronal differentiation promoted by these TFs.

TGIF2 also regulates various chromatin factors and histone modifiers, including *Arid1b*, *Arid4b*, and the histone methyltransferases/demethylases *Setbp1*, *Kdm1a*, and *Kdm7a*. Histone modifications, such as H3K36 methylation and H3K4 methylation in the context of bivalent marks, have been implicated in establishing epigenetically primed and “poised” transcriptional states^55^. Thus, TGIF2’s regulatory influence may extend beyond direct transcriptional repression, potentially contributing to neurogenic priming through additional epigenetic mechanisms.

Interactome analysis further determined factors that cooperate with TGIF2 to mediate repression, such as SIN3A and NURD repressor complexes. Functional assays using the TGIF2-KRAB and TGIF2-VP64 fusion proteins further reinforced its role as a transcriptional repressor, as shown before^20,32^. SIN3A, in particular, regulates diverse cellular processes such as cell cycle, differentiation, and development^56,57^, and has been implicated in neurological disorders such as intellectual disability, as well as cancer progression^59,60^, some of the previously described roles of TGIF2^19,20^. TGIF2 appears to guide the SIN3A complex to specific DNA targets, restricting the expression of primed genes and fine-tuning the transcriptional regulation of neurogenesis and neural differentiation. Additionally, we identified ARID4B, a component of the SIN3A complex, as a critical TGIF2 interactor as *Arid4b* KD abolished TGIF2 function. In mouse embryonic stem cells, *Arid4b* KD led to downregulation of differentiation programs of mesoderm and endoderm fate^61^. This is interesting in light of *Arid4b* also being a target of TGIF2 and hence reduced in expression by TGIF2. *Arid4b* KD in E12 cortex cells shows a trend of slight increase in Pax6+ NSCs compared to the control, although mild, but in the same direction of TGIF2 overexpression. Altogether, this indicates a negative feedback loop–TGIF2 interacts with ARID4B, and this complex represses the *Arid4b* transcript–as a molecular pathway regulating neural differentiation programs. Also, *Arid4b* KD was shown to increase globally H3K27me3 repressive histone marks^61^. The repressive histone mark H3K27me3 is particularly enriched at genes involved in neuronal maturation, serving as an epigenetic barrier during cortical development to ensure a protracted neurogenesis in human^62^. This resonates with TGIF2 overexpression phenotype that cells remain longer in progenitor state and a delayed neural differentiation.

It is worth noting that TGIF2’s function in the developing nervous system differs significantly from its role in other tissues. Unlike its reported interactions with SMAD proteins to regulate TGFβ target genes in other contexts, TGIF2 was not found to interact with SMAD proteins in this study. Additionally, while in many cancer cells TGIF2 promotes epithelial-mesenchymal transition (EMT), e.g. in lung adenocarcinoma (LUAD) cells^20^, it maintains the epithelial-like NSCs in the developing cortex as shown here. Therefore, TGIF2’s role in the nervous system exhibits significant mechanistic differences compared to cancer cells and endoderm-derived tissues, where it has been more extensively examined^22,26^. Most importantly, it was never characterized in priming and no major factors regulating neurogenic priming were previously known.

In summary, our findings establish TGIF2 as a master regulator of neurogenic priming and NSC fate across regions, using transcriptional repression to ensure the precise timing of cortical development.

## Methods

### RNA-seq and ATAC-seq libraries preparation

Wild type C57BL/6J embryos at E14 and E18 were used for the RNA sequencing experiments, with tissue of one litter/mother being pooled and considered one biological replicate. Brains were dissected in 1× HBSS (Gibco, cat. no. 14025) with 10 mM HEPES (Gibco, cat. no. 15630). Lateral cortex from the mediolateral to the cortex-LGE border, and LGE without overlying ventrolateral cortex, were dissected and centrifuged at 1000 rpm, 4 °C for five minutes. Dissection buffer was aspirated, and tissue was enzymatically dissociated with 1 ml of 0.05 % Trypsin/EDTA (Gibco, cat. no. 25300) for 15 minutes at 37 °C. Digestion was inhibited by adding 2 ml DMEM (Gibco, cat. no. 61965) with 10 % FBS (PAN Biotech, cat. no. P30-3302) and tissue was further mechanically dissociated with a fire-polished glass Pasteur pipette coated with DMEM + 10% FBS to obtain a single-cell suspension. The suspension was centrifuged at 1000 rpm, 4 °C for 5 minutes, the supernatant aspirated and the cells resuspended in 1× Staining Solution (1x HBSS, 1% Glucose, 1M HEPES, 1% FBS, 0.1% w/v NaN_3_, 1mM EDTA and DMEM-F12). The cell suspension was stained with the pre-absorbed antibody mCD133-PE at 1:500 dilution (Anti-Mouse-CD133-PE [13A4], eBioscience/Invitrogen, cat. no. 12-1331-82). A corresponding isotype control antibody (Mouse IgM-APC, Miltenyi Biotec, cat. no. 130-093-176) was added to an isotype control sample in the same dilution. Cells were incubated at 4 °C in the dark for 25 minutes, then DAPI (1:1000 dilution of 1 mg/ml stock; Sigma-Aldrich, cat. no. D9542) was added followed by another 5 minutes of incubation. To wash the cells, the suspension was filled up to 10 ml with PBS (Gibco, cat. no. 14190) and centrifuged at 1000 rpm, 4 °C for 5 minutes. Cells were resuspended in PBS and filtered through a cell strainer (pluriStrainer Mini 40 µm, PluriSelect, cat. no. 43-10040-60) into suitable sample tubes (Falcon™ Round Bottom Polypropylene Test Tubes with Cap, Falcon, cat. no. 352063).

Cells were sorted on a FACSAria™ III Cell Sorter (BD Biosciences) with FACSDiva software (version 6.1.3, BD Biosciences). To separate the populations the first gate was set to separate small debris (low FSC) and dead or damaged cells, which were DAPI+ (high 450/40 signal). The second gate was set to remove doublets or cell aggregates by FSC-area/FSC-width. The third gate separated the stained populations by the laser lines 582/15 for PE, with the gate set so that max. 0.1 % of the parent population in the isotype control was detected as single or double positive. Sorted cells were collected in PBS and centrifuged at 1000 rpm, 4 °C for 10 minutes. The supernatant was aspirated, and cells were immediately lysed in RNA extraction buffer. For the RNA-seq libraries, total RNA extraction was performed with the PicoPure™ RNA Isolation Kit (Applied Biosystems, cat. no. KIT0204) according to the manufacturer’s protocol with on-column DNase digestion (On-Column DNase I digestion set, Sigma-Aldrich, cat. no. DNASE70). RNA concentration and quality were evaluated on the Bioanalyzer (Model 2100, Agilent) using the RNA 6000 Pico Kit (Agilent, cat. no. 5067-1513) according to the manufacturer’s protocol. Samples with an RNA Integrity number (RIN) <8.0 were excluded from library preparation. First-strand cDNA was prepared from 2 ng RNA per sample with the SmartSeq v4 Ultra Low Input RNA Kit for Sequencing (TaKaRa/Clontech, cat. no. 634897) according to the manufacturer’s instructions. Number of amplification cycles for each sample was determined with a side qRT-PCR reaction performed after the first 4 amplification cycles to avoid over-amplification bias. With this, the number of required total amplification cycles for each sample corresponded to the cycle number at ¼ of the maximum fluorescence signal (Rn).

The amplified cDNA was purified using AMPure XP magnetic beads (Beckmann Coulter, cat. no. QT650) and quality and quantity analyzed by Bioanalyzer (High Sensitivity DNA Kit, Agilent, cat. no. 5067-4626) and Qubit Assay (Qubit™ dsDNA HS Assay Kit and tubes, Invitrogen, cat. nos. Q32854/Q32856). Purified cDNA was fragmented by ultrasonic shearing on the Covaris AFA S220 system using corresponding tubes (microtube AFA Fiber Pre-Slit Snap-Cap 6×16mm, Covaris, cat. no. 520045), resulting in approximately 200 bp – 500 bp long fragments that were purified by ethanol precipitation. Samples were evaluated again on the Bioanalyzer (HS DNA assay) before proceeding to the library preparation with the MicroPlex Library Preparation Kit v2 (Diagenode, cat. no. C05010014) according to the manufacturer’s instructions, using 10 ng of cDNA per sample. Following the library amplification, cDNA concentration was verified by Qubit assay and the libraries were purified over AMPure XP magnetic beads. Quality and quantity of these final libraries was evaluated by Bioanalyzer HS DNA assay and samples were multiplexed at 5nM each. Next generation sequencing was performed on an Illumina HiSeq 4000system with 100-bp paired-end deep sequencing.

For the ATAC-seq libraries, nuclei were isolated from 50,000 cells using a cell lysis buffer containing Tris-HCl 1M, NaCl 5M, MgCl2 1M, 10% NP40, 10% Tween-20 and 2% Digitonin. They were subsequently resuspended in transposition mixture containing the transposase enzyme, 2% digitonin and 10% Tween-20 and incubated for 30 minutes at 37°C. After the incubation the samples were immediately put on ice and DNA was purified with the MinElute Reaction Cleanup kit (Qiagen, #28204). The transposed DNA was PCR amplified with the NEBNext High-Fidelity 2x PCR Master Mix (NEB, #M0541S). The number of cycles was determined with a qRT-PCR using the SensiMix SYBR No-ROX 2x Master Mix (Bioline, #QT650) as the number of cycles that corresponds to ¼ of the maximum fluorescence. The amplified libraries were purified and the quality was assessed with a High Sensitivity DNA Chip (Agilent, #5067-4626). Size selection between 100bp and 600bp was performed with AMPure beads (BeckmannCoulter, #A63881) and libraries were pooled and sequenced on an Illumina HiSeq 4000system with 100-bp paired-end deep sequencing.

### RNA-seq analysis

The quality of sequencing data was analyzed with FastQC v0.11.4^63^ and adapter trimming was performed with cutadapt v1.11^64^. Reads were aligned with the mouse reference genome (mm39) using STAR v2.6.0a^65^. Afterwards, reads were deduplicated and gene expression was quantified with featureCounts v1.6.4^66^. The subsequent analysis was performed in R version 4.4.1^67^. Genes with less than 10 counts across all samples were excluded. The expression data was normalized and transformed using the vst function of DESeq2 v1.44.0^68^ for plotting and outliers’ analysis. To identify outliers, we performed a principal component analysis (PCA). Samples with a distance of more than 2.5 standard deviations from the mean in the first principal component were excluded (no outliers were detected). Differential expression (DE) analysis was performed using DESeq2 v1.44.0^68^. DE analysis for each comparison was done separately. We tested for DE with DESeq2 using the Wald test and reported the genes with a false discovery rate (FDR) below 1% as significant. Overrepresentation analysis for GO-Biological processes, Molecular pathways and Cellular compartment was done using ClusterProfiler v4.12.6. As background we used all genes on our dataset (22,125 genes). For all analyses, we used an FDR cutoff of 1% as significant threshold. Grouped semantic representation analysis was used to plot the significantly enriched GO terms. For this we used hierarchical clustering with the “Ward.D2” clustering method and Jaccard similarities. All data were plotted using the ggplot2 v3.5.1 package.

For identification of transcription factors, we used the GO term GO:0140110. For identification of chromatin remodelers, we used the GO terms: GO:0034724, GO:0031497, GO:0031498, GO:0034401, GO:0006338, GO:0016569, GO:0090202, GO:0070828, GO:0034728, GO:0006342.

### ATAC-seq analysis

FastQC was used to assess initial data quality. Reads were trimmed using *trim-galore* with parameter *--nextera* after contamination of Nextera Transposase Sequence was found in the reads. After trimming, reads were aligned to mm39 reference genome using *bwa-mem.* The *ATACseqQC* R-package tutorial was followed to assess data quality and to shift reads by 5bp as recommended^69^. For each individual sample, peaks were called using MACS3 with parameters *-f BAMPE -g mm -q 0.01*^70^. Differential openness of peaks between either time points per tissue or tissues per time point was identified using *DiffBind*with parameter *peakFormat=”narrow”* when loading the samples. *Homer* was used to find motifs in the resulting differentially open peaks. *Homer* was also used for labeling the differential or consensus peaks by genes in proximity. MonaLisa was used for motif enrichment analysis on differential or consensus peaks^71^. *ShinyGO 0.80*^72^ was used to assess pathway enrichment of the genes in proximity to peaks. Overlaps between peaks were identified by the function*subsetByOverlap.* From the 44 neurogenic fate determinants of Figure 1C, five had known binding motifs: *Atf3*, *Etv6*, *Mafk*, *Mycn* and *Tgif2*.

### Plasmids

TGIF2 cDNA isoforms plasmids were obtained from as a kind gift from previously described^26^. All plasmids for expression were cloned into a Gateway (Invitrogen) form of pCAG-IRES-GFP (kind gift of Paolo Malatesta) through pENTR1a vector. TGIF2 cDNA were amplified by PCR with primers containing triple FLAG sequence for inserting the FLAG tag at N-terminus of TGIF2 and cloned into the pCAG plasmid via Gibson Assembly. shRNA plasmids were designed using Invitrogen Block-iT RNA designer and ordered as oligos from Eurofins, then ligated to pENTR1a vector with a GFP reporter, which was finally cloned into a pCAG destination vector via Gateway LR clonase.

### Mice

The animals were housed in the Core Facility Animal Models (CAM), Biomedical Center (BMC), Faculty of Medicine, LMU Munich. They were maintained under specified pathogen-free conditions and housed in groups of 2-3 animals in individually ventilated cage systems with a 12 h/12 h light/dark cycle. C57BL/6J mice (Charles River Laboratories; Sulzfeld, Germany) were utilized for this study, and all animals undergoing in utero electroporation were females aged between 3 and 6 months. Embryonic day 0 (E0) was designated as the day of vaginal plug detection. Mice had free access to water and standard rodent chow (Altromin, 1310M). Experimental procedures were performed in accordance with animal welfare policies and approved by the Government of Upper Bavaria (Germany).

### Anesthesia

For surgical procedures, mice were anesthetized via intraperitoneal injection of a solution containing Fentanyl (0.05 mg/kg), Midazolam (5 mg/kg), and Medetomidine (0.5 mg/kg). Anesthesia was terminated with a subcutaneous injection of a solution comprising Buprenorphine (0.1 mg/kg), Atipamezole (2.5 mg/kg), and Flumazenil (0.5 mg/kg).

### *In Utero* Electroporation

Pregnant dams at E13 were anesthetized and operated on according to established procedures^44^. Briefly, endotoxin-free plasmids at 0.5 to 0.7 μg/μl, controlled for molar ratio across conditions, were diluted in 0.9% NaCl and mixed with FastGreen FCF dye. Subsequently, 1 μl of this mixture was injected into the lateral ventricle of embryos at E13 within anesthetized C57BL/6J mice. Embryonic brains were harvested at 3 days post-electroporation and fixed using 4% paraformaldehyde (PFA) in 1× PBS for durations of 4 hours. Analysis involved embryos obtained from at least two female mice, with quantification carried out on two to three coronal sections from three to five embryos.

### Cell culture

Cerebral cortices from C57BL/6J E12 mouse embryos were dissected in ice-cold Hanks’ balanced salt solution buffered with 10mM HEPES (both from Life Technologies). Cells were enzymatically dissociated with 0.05% Trypsin and mechanically triturated with a Pasteur pipette to obtain a single-cell suspension. These cells were then seeded in poly-d-lysine-coated coverslips in 24-well plates at 350,000 – 500,000 cells per well in DMEM-GlutaMAX supplemented with 10% FBS and 1% Pen/Strep and incubated at 37°C with 5% CO_2_. After 24 hours, 2% B27-supplemented DMEM-GlutaMAX with 1% Pen/Strep were added at 1:1 ratio. Three days or 7 days post transfection, cells were fixed with 4% PFA for 10 min at room temperature.

For transfection experiments, cells were plated and allowed to adhere for 2-3 hours before transfection with either 0.5 to 0.7 μg of plasmids controlled for molar ratio, or 25nM siRNA Tgif2 mouse (ON-TARGETplus SMARTpool) using Lipofectamine™ 2000 following the manufacturer’s guidelines (Invitrogen™). When shRNAs were co-transfected with GFP or TGIF2IR overexpression plasmids, equal molarity ratio was controlled.

### Immunohistochemistry and Immunocytochemistry

Sections underwent triple washes with 1× PBS at room temperature before being incubated overnight at 4°C with primary antibody in a blocking solution, composed of 10% Normal Goat Serum and 0.5% Triton-X100 in 1× PBS. Cells were first incubated in blocking solution for 1 hour at room temperature, followed by overnight incubation with primary antibody. After triple wash with 1× PBS at room temperature, cells and sections were stained with secondary antibodies diluted in blocking solution for 1 hour at room temperature. Nuclei were visualized using 0.5μg/ml 4,6-diamidino-2-phenylindole (DAPI, Sigma-Aldrich). Finally, immunostained sections and cells were examined using a Zeiss confocal microscope. The list of antibodies utilized in the experiments is provided for reference.

### scRNA-seq library prepration

36 hours after IUE, cortices were dissected in ice-cold Hanks’ balanced salt solution buffered with 10 mM HEPES (both from Life Technologies) under florescent microscope to enrich for electroporated region. The cells were dissociated to arrive at single cell suspension with Neural Tissue Dissociation Kit(P) (Milteny, #130-092-628) and red blood cell removal solution (Miltenyi, #130-094-183) following manufacture’s protocol. The cells were passed through a 40μm cell strainer and placed on ice for FACS to further isolate electroporate cells. FACS sorting was performed at a FACSAria III (BD Biosciences) in FACSFlow sheath fluid (BD Biosciences), with a nozzle diameter of 100 μm. Debris and aggregated cells were gated out by forward and side scatter, respectively. Single cells were selected by FSC-W/FSC-A. Gating for GFP fluorescence was done using non-electroporated cortices.

FAC-sorted cells were multiplexed using Cell Multiplexing Oligo Labeling and loaded onto 10X Chromium chip following Single Cell 3’ v3.1 (Dual Index) protocols with Feature Barcode technology for Cell Multiplexing (CG000388). The library was sequenced with one Novaseq 6000 S2 flowcell to reach 30,000 reads per cell for gene expression library and 5,000 reads per cell for multiplexing library, which was then aligned and demultiplexed using cellranger multi pipeline.

### scRNA-seq analysis

The analysis followed Scanpy’s^73^ tutorial, starting with preprocessing of raw sequencing data to filter out low-quality cells (counts per cell = 1100-33000, minimal genes per cell = 700) with high mitochondrial content (5% cutoff), followed by log transformation normalization. Dimensionality reduction using principal component analysis (PCA) and Uniform Manifold Approximation and Projection (UMAP) was performed to visualize cell-to-cell relationships. Leiden clustering identified distinct cell populations based on gene expression profiles, and marker genes were determined to characterize each cluster’s cell types. Maturation score included genes *Neurog2*, *Dcx*, *Tubb3*, *Elavl4*, *Map2*, *Stmn2*, *Rbfox3*, *Syt1*, *Nefl*, *Syn1*, *Syp*, *Camk2a*, Bsn. DE between TGIF2IRwt and GFP was analyzed using built-in “rank genes” function in Scanpy with Wilcoxon rank-sum test, and associated GO term was analyzed using Shiny GO 0.80^72^. DE between TGIF2IRmut and GFP was analyzed using pseudobulk and DESeq2 v1.44.0^68^ to be comparable to the bulk Cut&Run. CellRank analysis based on RNA velocity was conducted followed CellRank’s tutorial^28,29^.

### Cleavage under targets and release using nuclease (CUT&RUN) and library preparation

Electroporated embryos underwent the same procedure as described in scRNA-seq section until before FACS. Cut&Run was performed using CUT&RUN assay kit (Cell Signaling Technologies, 86652) according to the manufacturer’s instructions. Briefly, 250,000 cells per reaction were collected and bound to Concanavalin A Magnetic beads. Cells were permeabilized and incubated with 1 μg of primary antibody against FLAG (DYKDDDDK Tag (D6W5B), rabbit, Cell Signalling) per sample overnight at 4°C. The rabbit (DA1E) mAb IgG XP® Isotype Control antibody was used as IgG control. Subsequently, cells were incubated with pAG-MNase for 1 h at 4°C. pAG-MNase was activated by adding calcium chloride and incubation at 4°C for 30 minutes. Stop buffer (Cell Signaling Technologies) was added to each sample to stop the reaction. DNA was purified using phenol/chloroform extraction and ethanol precipitation as described in the manufacturer’s protocol.

DNA sequencing libraries were generated using the SimpleChIP® ChIP-seq DNA Library Prep Kit for Illumina (Cell Signaling Technologies, 56795) and SimpleChIP® ChIP-seq Multiplex Oligos for Illumina® (Dual Index Primers, Cell Signaling Technologies, 46538) following the manufacturer’s instructions specifically for CUT&RUN Assay kit protocol. Briefly, 5ng of DNA was used for all CUT&RUN and IgG control samples. DNA ends were ligated with adaptors and amplified using PCR and Dual Index primers for Illumina® (Cell Signaling Technologies, 47538). All clean-up steps were performed with 1.1× volume of SPRIselect® beads to increase the capture of smaller DNA fragments. Generated libraries were pooled and sequenced using 2 × 75 bp paired-end sequencing strategy on an Illumina® NextSeq550 sequencer.

### Cut&Run analysis

Sequenced reads were aligned to the mm39 genome using Bowtie2^74^. Peak calling was performed using the MACS3 pipeline^70^ with corresponding IgG control bam files, using q value 0.01, and minimal fragment length 100. An enrichment heatmap of the peaks was produced using deepTools’s computeMatrix function^75^ on Galaxy platform^76^. FIMO motif scanning was conducted on MEME Suite website using bed file of identified peaks^77^. The peaks were analyzed for genomic distribution with ChIPpeakAnno^78^ and annotated using GREAT for single nearest gene within 250kb^30^. GO term enrichment analysis with ShinyGO 0.80 was conducted on the annotated genes^72^.

### Gene regulatory dynamic analysis

RegVelo is an end-to-end deep generative model designed to infer cellular dynamics through coupled splicing dynamics and gene regulation^31^. It requires users to define the prior gene regulatory network and allows the model to refine this network by improving the reconstruction of observed gene expression. Using a bulk ATAC-seq dataset, we followed CellOracle’s tutorial^79^. First, we identified transcription start sites (TSS) using the get_tss_info function, which annotates each peak with its corresponding gene. Next, we scanned transcription factor (TF) binding motifs in these peak regions using the tfi.scan function with an FPR of 0.02. Subsequently, we filtered motifs using the filter_motifs_by_score function with a threshold of 10. Finally, we replaced the bulk ATAC-seq-derived TGIF2 targets with CUT&RUN-inferred target genes and incorporated this prior GRN for downstream RegVelo analysis.

We trained the RegVelo model with default parameters. To mimic overexpression effects, we manually perturbed the inferred gene regulation by multiplying TGIF2 downstream regulation weights by a specific factor to amplify the regulatory effects of TGIF2. We used four different values [0, 50, 100, 150] and employed RegVelo to predict the depletion scores^31^ for defined terminal states, including NSCs, UL neurons, and DL neurons. RegVelo-inferred GRN targets were used for downstream gene functional analysis. We curated all negatively regulated genes inferred by RegVelo and applied the clusterProfiler package to perform GO enrichment analysis.

### Coimmunoprecipitation

For interactome analysis, P19 cells were seeded in 10cm dishes for transfection when the cells reached 50% confluency. After 48 hours, cells were scraped on ice and lysed in non-denaturing lysis buffer (20mM Tris-HCl, pH 8.0, 137mM NaCl, 1% Nonidet P-40, 2mM EDTA) containing cOmplete proteinase inhibitor. Lysates were incubated with DYKDDDDK Tag (D6W5B) FLAG rabbit antibody (Cell Signaling) for 1 hour, followed by addition of Protein G Dynabeads for an additional 2 hours at 4°C with rotation. Following three washes with wash buffer (10mM Tris, pH 7.4, 1mM EDTA, 150mM NaCl, 1% Nonidet P-40), the immunoprecipitated lysates were boiled in 1× Laemmli buffer and subsequently stored at –80°C until mass spectrometry analysis.

### Mass spectrometry

The interactome samples were digested using a modified FASP procedure as described^80,81^. Digested peptides were measured on a QExactive HF X mass spectrometer (Thermo Scientific) online coupled to an Ultimate 300 nano-RSLC (Thermo Scientific) as described^82^.Generated raw files were quantitatively analyzed in the MaxQuant software^83^ (MPI Martinsried, version 2.4.9.0), applying default settings and a minimum LFQ ratio count of 1, quantification on unique peptides with matching between runs for LFQ quantification^84^. Searches for peptide identifications were performed in the integrated search engine Andromeda^85^ with default settings, using the canonical SwissProt Mouse protein database including the described TGIF2 sequences. Results were filtered for contaminant hits, reverse hits and “only identified by site” hits. LFQ intensity values in the filtered proteingroups list were used for enrichment ratio calculations.

### Western Blot

P19 cells transfected with various shRNA constructs were lysed with RIPA buffer and the proteins were extracted by centrifugation at 13,000 x g for 15 minutes at 4°C. 30 μg protein per sample was diluted to the desired concentration in 1× Laemmli Buffer and boiled at 95°C for 5 min. Gel electrophoresis was conducted using 12.5% polyacrylamide SDS gels, followed by transfer to nitrocellulose membranes. For immunodetection, membranes were initially blocked with 5% nonfat dry milk in TBS-T (Tris-buffered saline/0.1% Tween20, pH 7.4) for either 1 hour at room temperature or overnight at 4°C, and then incubated overnight with primary antibodies (ARID4B, Bethyl Laboratories, 1:2000) diluted in 1% nonfat dry milk in TBS/T. The following day, the membranes were incubated with HRP-coupled secondary antibodies diluted in 1% nonfat dry milk in TBS-T. Finally, the signal was visualized using the ECL method with the ChemiDoc™ instrument from Biorad.

### Statistical analysis

The statistical tests were performed using GraphPad Prism 9. If the data passed the Shapiro-Wilk normality test, and F test (two conditions) or Barlett’s test (three or more conditions) for equal variance, they were subject to either unpaired t-tests when there were two conditions, or ordinary ANOVA with Tukey’s multiple comparisons test when there were three or more conditions. If the data passed the normality test but not equal variance, they were subject to Welch t-test when there are two conditions, or Brown-Forsythe and Welch ANOVA tests with Dunnett’s T3 multiple comparisons test when there were three or more conditions. If the data did not pass the normality test, they were subject to Mann-Whitney test when there were two conditions, or Kruskal-Wallis ANOVA with Dunn’s multiple comparisons test when there were three or more conditions.

## Acknowledgements

We are particularly grateful to Hyung-Seo Kang, Arie Geerlof and Michael Sattler for excellent input on the structure of TGIF2 including the mutations. Special thanks to Tatiana Simon-Ebert, Martina Buerkle, Paulina Chlebik for excellent technical support. We acknowledge the Core Facility Bioinformatics at the Biomedical Center, LMU Munich, especially Tobias Straub for consultation on Cut&Run analysis and HPC server usage, and then to Core Facility Flow Cytometry (Benjamin Tast and Roqayeh Noori) at the Biomedical Center, LMU Munich for FACS of cells for the scRNA-seq experiments. Sequencing of the bulk libraries was performed at the NGS facility, Institute of Human Genetics, Helmholtz Center Munich. Sequencing of the rest of the experiments was performed at the Next Generation Sequencing (NGS) Core Facility at the Institute of Human Genetics in Bonn. We are also very grateful to Giacomo Masserdotti and Stefan Stricker for excellent comments on the manuscript.

This study was supported by the advanced ERC Grant Neurocentro (885382 to M.G.) and the European Union’s Horizon 2020 research and innovation program under grant agreement no. 874758 (NSC Reconstruct to M. G.), as well as the German Research Foundation TRR274 (no. 408885537, M. G.), FOR2879/2 (no. 405358801, M. G.) and SyNergy (EXC2145/Project-ID 390857198, to M.G.). Parts of this work were supported by a New Frontiers in Research Fund Transformation grant to M. Götz, funded through three Canadian federal funding agencies (CIHR, NSERC, and SSHRC).

## Contributions

M.G. conceived the project and together with Y.L. designed the study. Y.L. performed experiments and data analysis, including cloning, IUE experiments, E12 transfection experiments, imaging, data analyses, Cut&Run experiment and analysis, scRNA-seq experiment and analysis, IP experiment. F.V. acquired bulk RNA- and ATAC-seq data. A.K analyzed bulk RNA-seq data and M.R. analysed ATAC-seq data and priming data. W.W. performed RegVelo analysis and F.T. advised on the analysis. J.M-P. and S.H. performed proteomics of IP samples. Y.L., A.K., M.R., and M.G. wrote the manuscript, with input from all co-authors. M.G. provided all the funding.

---

**Figure S1.**
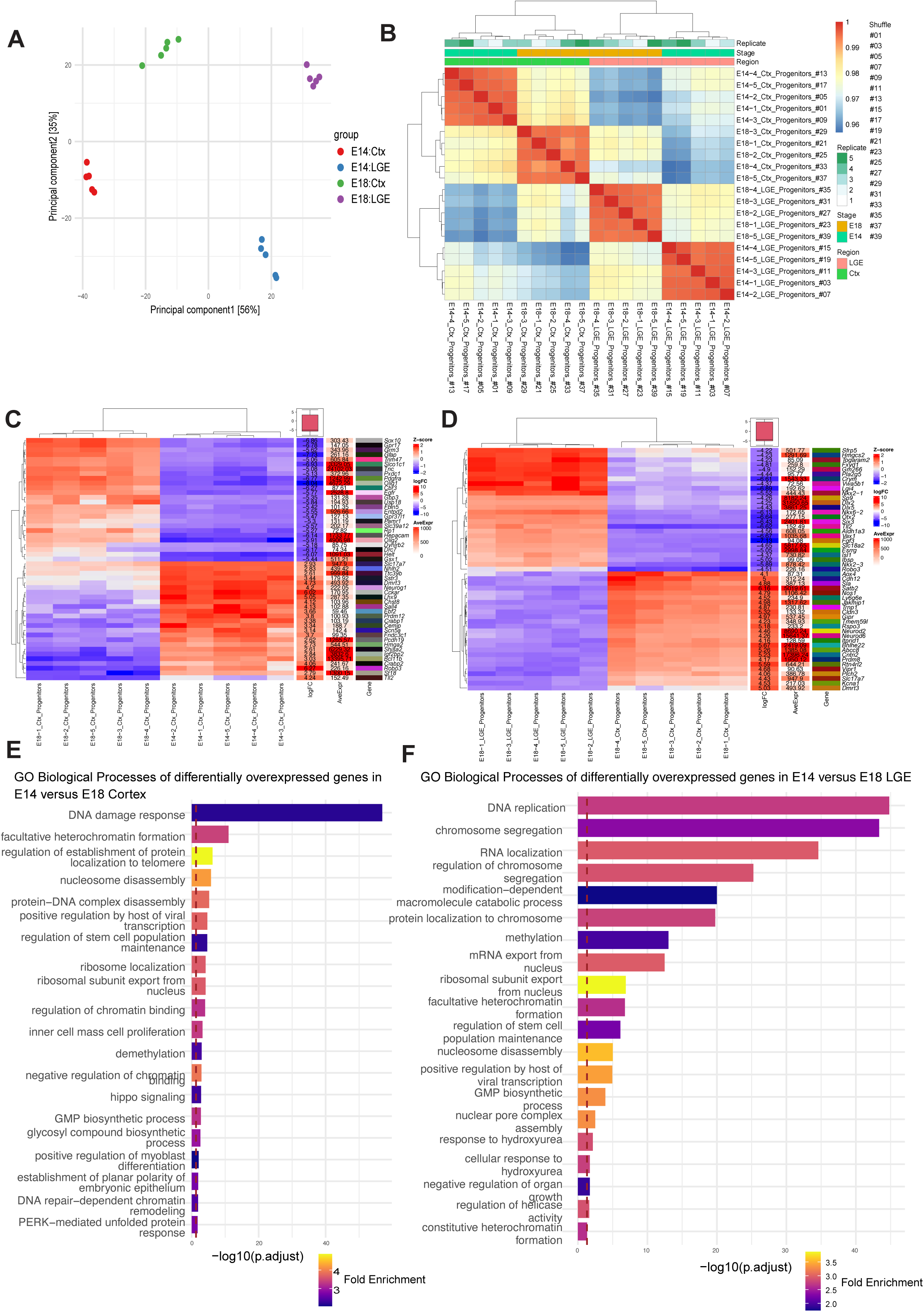
RNA-seq analysis of Radial Glial cells isolated from cerebral cortex and LGE at E14 and E18. (A) Principal component analysis of the RNA-seq data with region marked with different shapes and stage marked with different colours. Ctx: Cortex; LGE: Lateral Ganglionic Eminence; E: Embryonic (B) Heatmap of samples clustered according to different parameters of the dataset. (C) Heatmap of the top 25 differentially up- or down-regulated genes in the cortex at E14 versus E18. FC: fold change; AveExpr: average expression (D) Heatmap of the top 25 differentially up- or down-regulated genes in the LGE at E14 versus E18. FC: fold change; AveExpr: average expression (F) GO terms associated with biological processes, showing top 2 terms each from 10 clusters of semantic space analysis, taken from genes upregulated at E14 versus E18 cortex. (G) GO terms associated with biological processes, showing top 2 terms each from 10 clusters of semantic space analysis, taken from genes upregulated at E14 versus E18 LGE.

**Figure S2.**
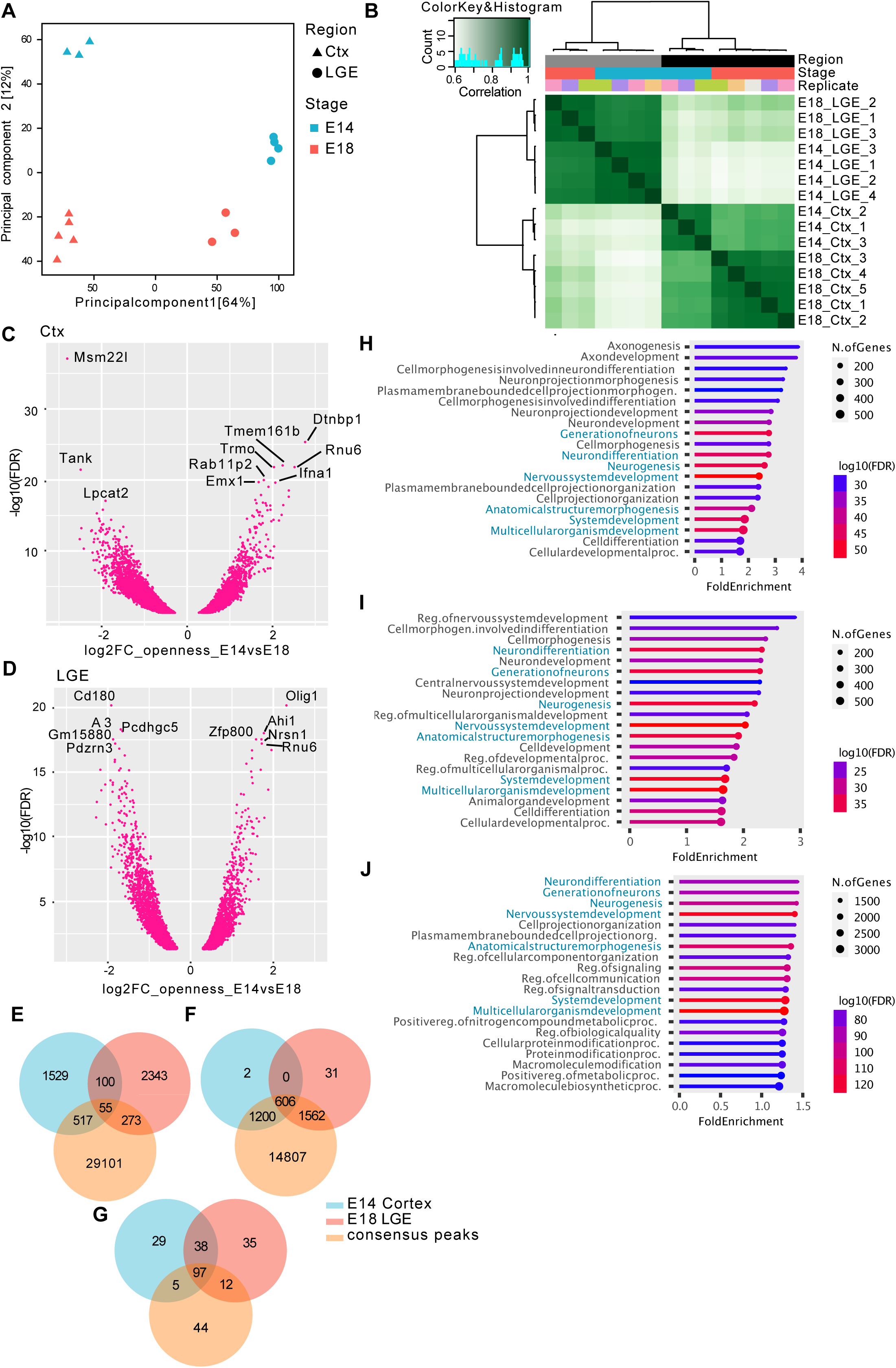
ATAC-seq analysis of Radial Glial cells isolated from cerebral cortex and LGE at E14 and E18. (A) Principal component analysis of the ATAC-seq data with region marked with different shapes and stage marked with different colours. Ctx: Cortex; LGE: Lateral Ganglionic Eminence; E: Embryonic (B) Heatmap of the ATAC-seq samples clustered based on chromatin openness and annotated based on different parameters of the dataset. (C) Volcano plot of significantly differentially open peaks in E14 versus E18 in Cortex (FDR < 0.05). Top 10 differential peaks are labeled by the nearest gene in their proximity. Ctx: Cortex; LGE: Lateral Ganglionic Eminence; E: Embryonic (D) Volcano plot of significantly differentially open peaks in E14 versusE18 in LGE (FDR < 0.05). Top 10 differential peaks are labeled by nearest gene in their proximity. Ctx: Cortex; LGE: Lateral Ganglionic Eminence; E: Embryonic (E) Venn diagram depicting overlap of differentially open peaks in E14 versus E18 Cortex, differentially open peaks in LGE versus Cortex E18, and the non-differential (consensus) peaks between Cortex and LGE in E14. Left: overlap of the peaks, middle: overlap of the genes in proximity to peaks, right: overlap of the motifs associated to open peaks. (F) Venn diagram depicting overlap of genes in proximity to differentially open peaks in E14 versus E18 Cortex, differentially open peaks in LGE versus Cortex E18, and the non-differential (consensus) peaks between Cortex and LGE in E14. (G) Venn diagram depicting overlap of motifs enriched in differentially open peaks in E14 versus E18 Cortex, differentially open peaks in LGE versus Cortex E18, and the non-differential (consensus) peaks between Cortex and LGE in E14. (H) Barplot showing gene ontology terms enriched in genes in proximity to differentially open peaks between E14 and E18 in Cortex. (I) Barplot showing gene ontology terms enriched in genes in proximity to differentially open peaks between LGE and Cortex in E18. (J) Barplot showing gene ontology terms enriched in genes in proximity to non-differentially open (consensus) peaks between Cortex and LGE in E14. Terms that are shared between all 3 comparisons are highlighted in blue.

**Figure S3.**
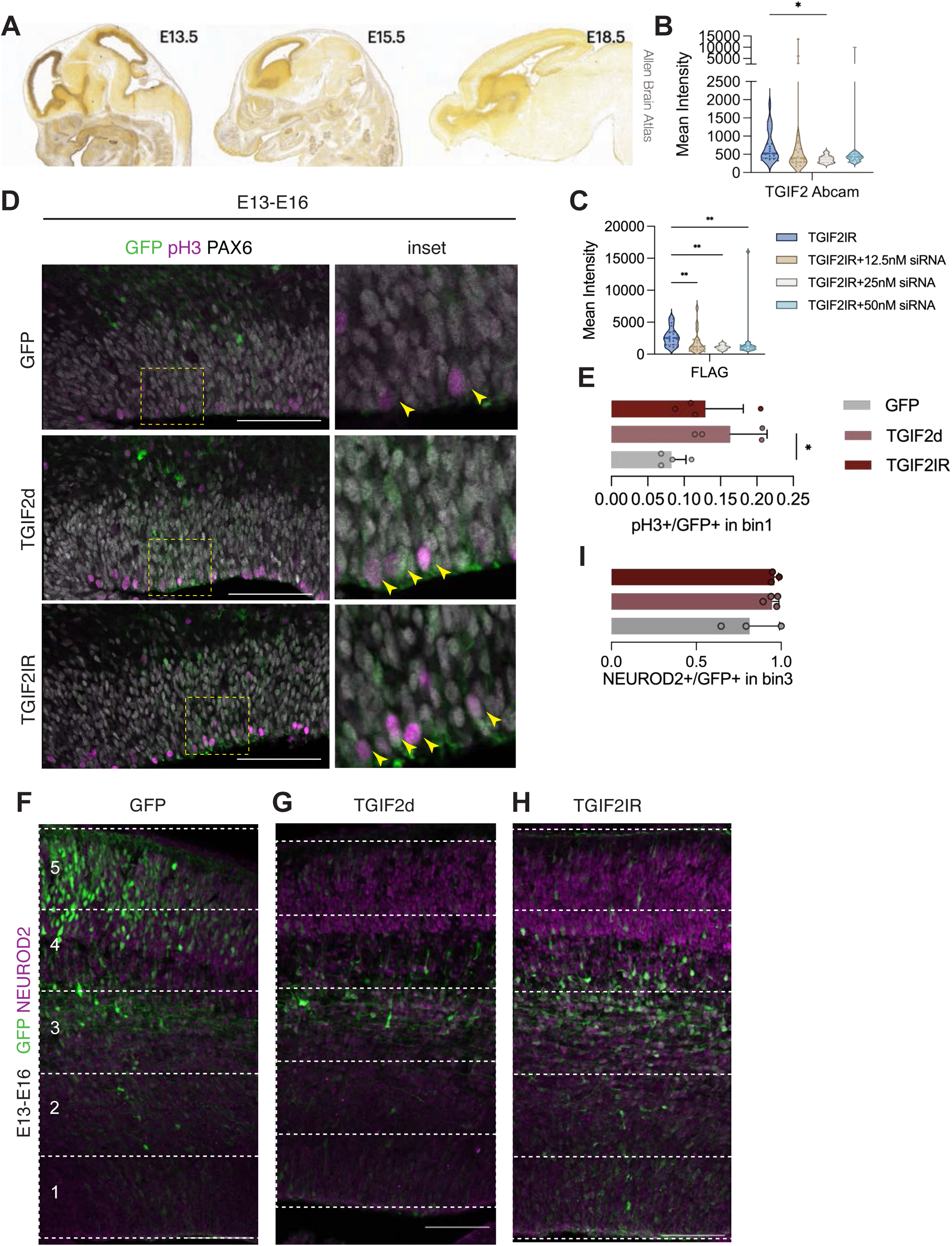
TGIF2 expression during development and immunostaining analysis after overexpression *in vivo*. (A) ISH data of mouse TGIF2 across different developmental timepoints, excerpts from Allen Brain Atlas^41^. (B-C) Violin plots showing quantification of knockdown efficiency titration with siRNA pool in final concentrations, together with TGIF2IR overexpression, measured by mean intensity in the channel of TGIF2 Abcam antibody (B) and the channel of FLAG antibody (C). N=8-21 cells measured with DAPI mask. Kruskal-Wallis ANOVA with Dunn’s multiple comparisons test. (D) Representative images and their insets of *in utero* electroporated cortices from different conditions (GFP+) immunostained with pH3 and PAX6. Arrowheads indicate pH3+/GFP+ cells. (E) Quantification of pH3+/GFP+ cells in bin 1, mean±SD. N=4 embryos from at least 2 mothers. Ordinary one-way ANOVA with Dunnett’s multiple comparison’s test. (F-H) Representative images showing cortices 3 days post electroporation with different conditions in GFP, co-stained with NEUROD2. Dashed lines indicate the 5 equal bins. Scale bar: 100μm (I) Quantification of NEUROD2+/GFP+ cells in bin 3, mean±SD. N=3-4 embryos from at least 2 mothers. Ordinary one-way ANOVA with Dunnett’s multiple comparison’s test. Only significant result is shown.

**Figure S4.**
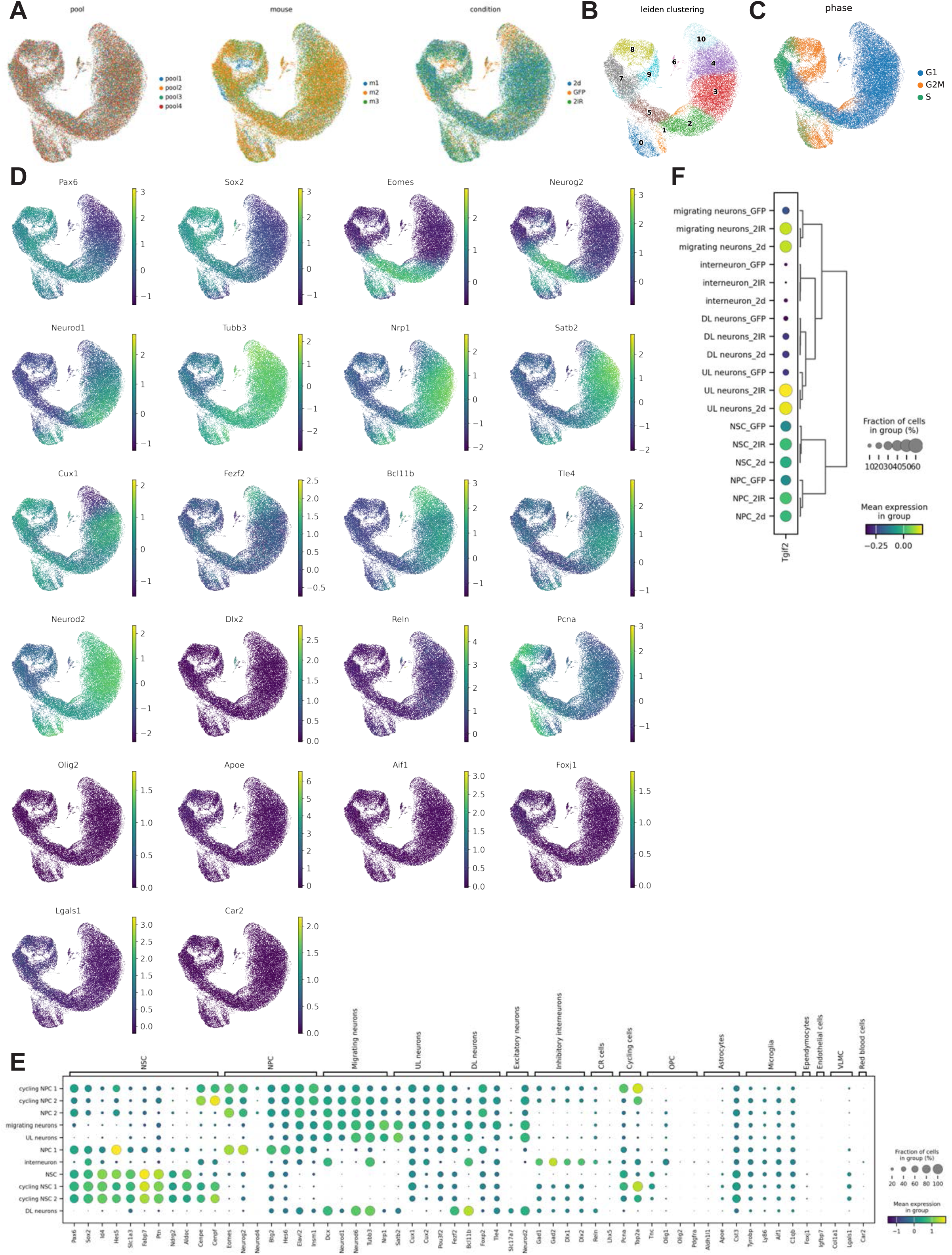
Leiden clustering and marker gene expression in the scRNA-seq data. (A) UMAP projection of cells grouped by pool, mouse, and condition. (B) Leiden clustering with UMAP projection. (C) Scatter plot of cell cycle phase from cell cycle marker gene expression. (D-E) UMAP scatterplot and dot plot of marker gene expression across cell clusters. (E) Tgif2 expression levels across different cell types and conditions.

**Figure S5.**
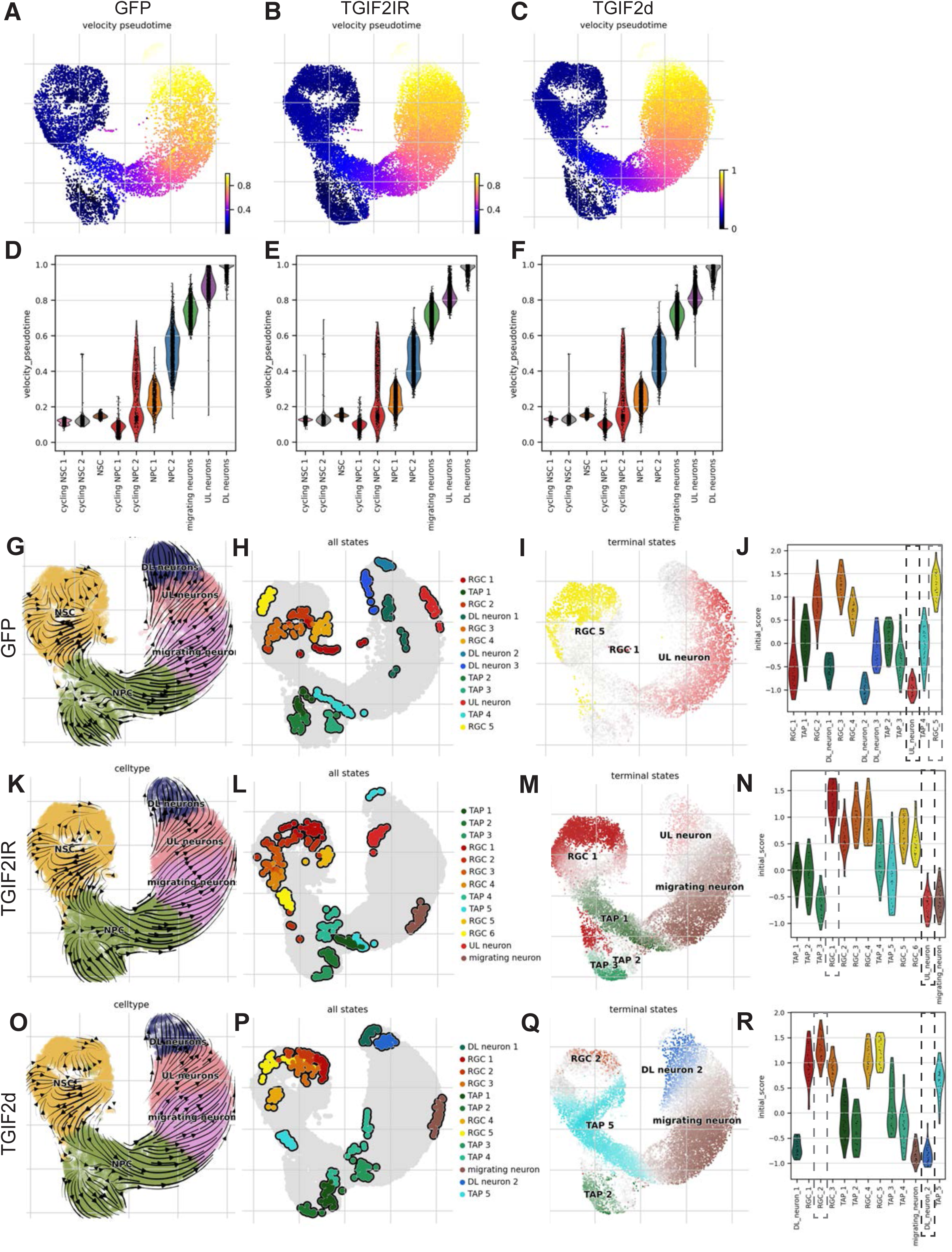
Velocity pseudotime and CellRank procedures. (A-C) UMAP of velocity pseudotime across 3 conditions. (D-F) Violin plots of velocity pseudotime across cell types in different conditions. (G, K, O) UMAP representation of RNA velocity^27^. (H, L, P) Macrostates predicted by CellRank. RGC: radial glial cells, or NSCS; TAP: transit amplifying progenitors, or NPCs. (I, M, Q) Terminal states predicted by CellRank^28,29^. (J, N, R) Violin plots of initial score (Fabp7, Pax6, Sox2 expression) of macrostates predicted by CellRank^28,29^.

**Figure S6.**
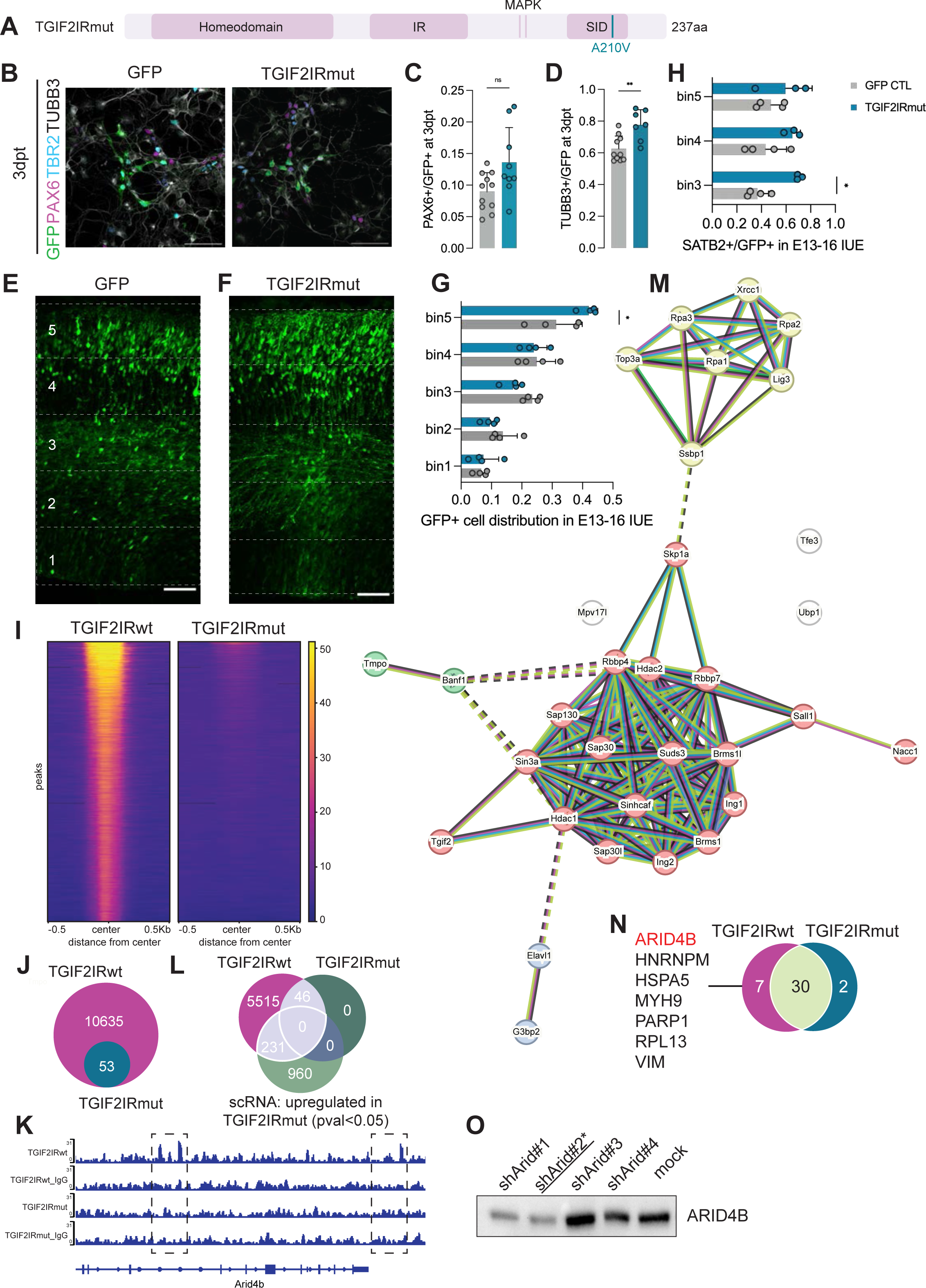
Mutation in SID of TGIF2 abrogates its function, some interactions and binding to target sites. (A) Schematic drawing of TGIF2IRmut construct. (B) Representative images showing E12 primary cortex cells cultures transfected with GFP, same as in Figure 6E, and TGIF2IRmut, co-stained with PAX6, TBR2, and TUBB3. Scale bar: 50μm. (C-D) Quantification of PAX6+/GFP+ and TUBB3+/GFP+ in transfected E12 cortex cell culture at 3dpt with different conditions as indicated in the legends, mean±SD. N= 7-11 pools of embryos. Mann-Whitney test. (E-F) Representative images showing cortices 3 days post IUE with GFP (E) and TGIF2IRmut (F). Dashed lines indicate the 5 equal bins. Scale bar: 100μm. (G) Quantification of GFP+ distribution in each bin, mean±SD. N=4 embryos from at least 2 mothers. Ordinary two-way ANOVA with Sidak’s multiple comparisons test, with a single pooled variance. (H) Quantification of SATB2+/GFP+ in each bin from embryos *in utero* electroporated with GFP or TGIF2IRmut. Ordinary two-way ANOVA with Sidak’s multiple comparisons test, with a single pooled variance. (I) Enrichment heatmap of TGIF2IRwt and TGIF2IRmut, centered on the middle of the peaks. (J) Venn diagram showing the overlapped *peaks* between TGIF2IRwt and TGIF2IRmut from Cut&Run analysis. (K) Peak examples with bigwig profile of TGIF2IRwt and TGIF2IRmut, with their corresponding IgG control at the gene locus of *Arid4b*. Dashed lines circle the peaks. (L) Venn diagram showing the overlapped *genes* between TGIF2IRwt and TGIF2IRmut from Cut&Run analysis, as well as genes upregulated in TGIF2IRmut compared to GFP control from its scRNA-seq DE analysis (log2fc>0, pval <0.05). (M) STRING analysis of interactors of TGIF2IRmut with LFQ intensity more than 3-fold compared to GFP control. (N) Venn diagram comparing the interactors of TGIF2IRwt and TGIF2IRmut with LFQ intensity more than 3-fold compared to GFP control. The interactors lost in TGIF2IRmut are listed. ARID4B, which belongs to SIN3A complex, is marked in red. (O) Western blot of P19 cells transfected with candidate shRNAs for Arid4b KD, including a mock transfection control. Blotted bands show canonical ARID4B protein. shRNA#2 (against open reading frame of *Arid4b*) with the most efficient KD was used for further experiments.

